# AIRUS: a simple workflow for AI-assisted exploration of scientific data

**DOI:** 10.1101/2025.02.23.639768

**Authors:** Kenneth D. Harris

## Abstract

The development “reasoning” large language models (LLMs) seems poised to transform data analysis in all fields of science. This note describes a simple, iterative workflow termed AIRUS (AI Research Under Supervision), that allows working scientists, including those who lack extensive AI or programming expertise, to immediately start taking advantage of these capabilities. In the proposed workflow, a researcher uses an LLM to generate hypotheses, produce code, interpret results, and refine its approach in repeated cycles, only intervening when necessary. This can be run by simple cut-and-paste of code and results between web-based LLMs and an online Jupyter notebook. We demonstrate the workflow with two examples: one analysis of synthetic data, and one analysis of data from the International Brain Lab.

## Introduction

The development of LLM “reasoning models” such as openAI’s o1, DeepSeek-R1, Gemini 2.0 Flash Thinking, and Grok 3 promises to transform scientific research. Not only are these models fast and accurate at writing computer code, they are increasingly capable of scientific reasoning, data interpretation, interpretating data visualizations, and hypothesis generation. Research toward automated scientific discovery has shown great promise recently ^1–12^, but does not always translate to practical actions working scientists can immediately take to increase their productivity. However, the capabilities of current LLMs are already sufficient to transform the daily practice of scientific research. This note describes a simple workflow that allows working scientists, including those with no background in AI systems or LLMs, to immediately start taking advantage of this.

The workflow is an aid for exploratory data analysis. After a scientist has collected a dataset, the first task is usually to explore it by iteratively making plots and computing statistics, both to look for potential artifacts, and to define and refine hypotheses for structure in the data. At the time of writing, LLMs are very good at coding, but their scientific domain knowledge and intuition in most fields is still behind that of most working scientists. A productive way forward is therefore “research under supervision”: the scientist gives the LLM a description of the problem, including as much specialist domain-knowledge as possible, then allows it to work independently, but stands ready to correct any mistakes or intervene if it enters a “blind alley”. At the very least, this approach provides the scientist a substantial time saving due to the LLM’s fast code-writing abilities. At best, it can provide the scientist with new ideas they would not have thought of otherwise.

### The AIRUS workflow

The proposed workflow is outlined in Figure 1. I make no particular claim of originality: something like this is probably already used by many scientists. Nevertheless, I hope that writing this note might speed adoption of this or similar approaches by a wider community. The proposed workflow could run through an API, but can also be used with commercial or free chatbot services, simply by cutting and pasting code and results between the chatbot and an online Jupyter notebook.

**Figure 1:**
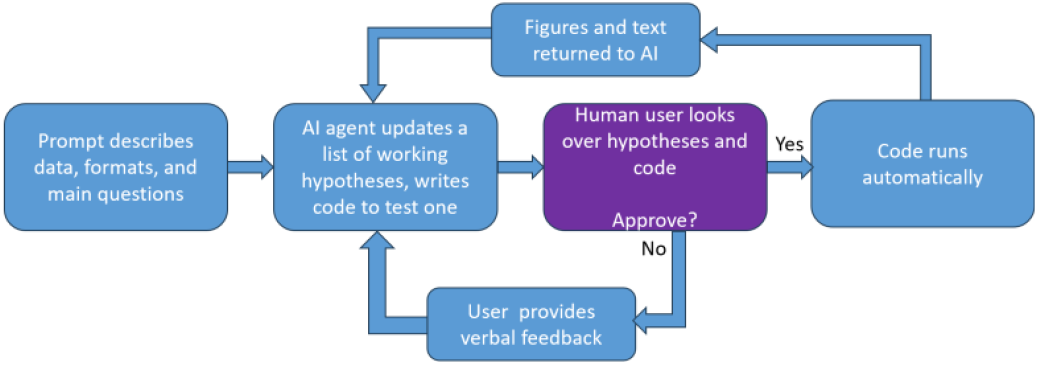
the AIRUS (AI Research Under Supervision) workflow.

To begin an analysis, the user crafts a prompt that describes to the LLM the iterative workflow that will be used, as well as describing the scientific question, the data to be analyzed, and the goals of the analysis. (Two example prompts are given in Appendix 1.) The prompt contains a specific description of the data to be analyzed (including array sizes), and as much domain-specific knowledge as possible: for example which statistical tests are likely to be useful and which invalid; which libraries to use or not to use; where experimental artifacts are likely to occur; and which kinds of reasoning are likely to be productive in this work (e.g. exploratory plots vs. significance tests). These are essentially the same kind of instructions and intuitions one would try to impart to a scientist new to a specific domain. The user sets up a Jupyter notebook system, and loads the data to be analyzed into memory. Note that **it may be unwise to run AI-generated code on computers with write access to your local filesystem**, so may be better to use a web-hosted Jupyter service such as Google Colab.

The workflow then proceeds in a series of iterations. On each iteration, the LLM summarizes its working hypotheses, selects one to test, and writes code to test it. The user reads this, and if happy, executes it and returns the output (text and graphics) to the LLM. If the user spots a problem, or notices something in previous results that the LLM has missed, this is added as free text input. The next iteration then starts. If necessary, one can define “phases” of analysis such as exploratory and confirmatory, and stopping criteria, in the initial prompt.

Better results are obtained if one gives the LLM a metric of success: for example a mean-square error metric or correlation to be optimized. Informal experimentation with different prompts suggests that giving such a metric results in faster convergence to good performance, presumably because it focuses the LLM on improving its performance over iterations.

Informal experimentation also suggests several other prompt features that improve performance: describing the exact workflow to the LLM; requesting a list working hypotheses on each iteration; and by encouraging graphical exploratory analysis, whose interpretation requires using vision-capable LLMs, that are able to interpret scatter plots and other exploratory graphical analyses.

Two examples of analyses run with this system are presented below.

### Example 1: synthetic data

The first example contains synthetic data. The aim is to predict a scalar variable *y* from a 3-dimensional multivariate *x*. The data are generated by sampling *x* from a standard uncorrelated Gaussian, and defining 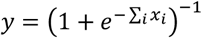. This is a logistic function, but the problem is not standard logistic regression, which would have a binary output. The prompt in Appendix 1a was given to OpenAI’s o1, the resulting code was run in Google Colab, and the resulting graphics and text produced were manually copied back to o1.

The workflow took 6 iterations to find the correct answer, with no human intervention. It first fit a univariate linear regression model to each variable independently; then found the fit was improved by multivariate linear regression, but noted that the residuals were structured suggesting nonlinearity. It then tried quadratic terms and cross-terms but found these did not markedly improve the fit. Finally, it noticed that a predicted-vs-actual plot (which it was not specifically instructed to make) had a sigmoid shape, and also that *y* lay in the range [0,1], so decided to apply a logit transform to *y*, which enabled the correct answer to be found.

This example illustrates the importance of three features of the workflow. First, defining a numerical metric of success (in this case mean-square error) enabled the LLM to understand that quadratic terms were of little benefit. Second, the key insight towards finding the logistic function came from visual analysis of an exploratory plot. An explicit instruction to produce such plots (and the use of a vision-capable LLM) was necessary for this to occur. Finally, the prompt contained information specific to the problem: in this case, an instruction not to use machine-learning libraries but to focus on simple numpy computations. In earlier attempts at the problem, without this instruction, the LLM obtained good prediction scores using approaches such as random forests, but did not discover the explicit formula.

### Example 2: relating neuronal autocorrelograms to spike waveforms

The second example focuses on a problem from neuroscience: relating the temporal structure of spike trains, as assessed by their autocorrelogram, to their spike waveforms. This example was chosen as it illustrates the kind of scientific question that could be accelerated by AI-assisted data exploration. The brain contains multiple types of neuron, spread over many areas, which may differ both in the temporal structure of their activity, and in the shape of the extracellular voltage waveforms (“spikes”) produced when they fire. The International Brain Lab ^13^ has performed recordings covering the whole mouse brain, which provide an opportunity to answer the question. In this initial test, we only analyzed one recording from the dataset.

Again, the analysis was performed by manually pasting code and results between o1 and Google Colab. While o1 has good overall knowledge of neuroscience, it was necessary to include specific domain knowledge in the prompt: for example, the fact that spike amplitude can be affected by distance of the neuron from the electrode, so waveforms should be normalized; and that the ACG range most likely to be useful was -200 to +200 ms. Specific instructions on how to use IBL’s quality control metrics to exclude low-quality units were provided, as was encouragement to start with graphical exploratory methods such as PCA and UMAP, and instructions to display ACGs and waveforms of example neurons. The latter was as much for the benefit of the human user as for the AI itself.

The workflow ran largely, but not completely, without human intervention. As instructed, the LLM started by visualizing principal component analysis of the ACG and waveforms. It also found (without explicit instruction) that the PCs differed between the recorded brain regions, and found a modest correlation between ACG and waveform PCs. It tried UMAP but did not find this separated the points from different brain regions. It then tried canonical correlation analysis (CCA) between PCA projections of the ACG and waveform, and found a stronger correlation. It tried some specific metrics (burst indices, half-widths, peak ratios, etc) but these did not come close to the correlation strength found by CCA.

At this point, the user intervened to say that running CCA on high-dimensional data can produce spurious correlations, and to suggest using cross-validation. The LLM performed this analysis, which verified the correlation. The user then requested additional plots, which led to a hypothesis that the CCA correlation represented in part a relationship between burst propensity and the size of the secondary positivity in spike waveform. The LLM confirmed this relationship, which to my knowledge was previously undescribed.

## Conclusion

A simple workflow of AI research under supervision was found to be useful for exploratory data analysis. The workflow and prompting strategy is simple, but even at this early stage, o1’s performance was roughly equivalent to that expected of a novice human researcher, but many times faster. This performance is likely to improve substantially in the near future: both by improvements in prompting strategies, and by improvements in the reasoning capabilities and domain knowledge of future LLMs themselves.

Informal experimentation suggested several prompt features that can improve performance. First, it helps to give the LLM a quantitative metric of success. This does not need to be explicitly implemented in code: instead it suffices to write a verbal instruction in the prompt.

Second, vision-capable models can make substantial progress by interpreting exploratory plots – but the LLM needs to be explicitly instructed to make these plots. Finally, providing as much domain-specific information as possible, including which approaches not to use, seems to improve performance.

Informally, it appears that LLMs tend to make similar mistakes in data analysis to novice human scientists: becoming fixated on particular approaches without considering alternatives; failing to explore how results depend on arbitrary analysis parameter values; rushing into confirmatory analyses and significance tests without first exploring the data graphically; and giving up too early after a single negative result when alternatives have not yet been tried. It is likely that these problems can be addressed with improved prompting. Nevertheless, even the simple approach tested here may already be enough to greatly accelerate progress in data analysis across scientific domains.

## Appendix 1a: prompt for synthetic data example

The text below was used as the initial prompt for the analysis of synthetic data. A full dump of the session (code and results) can be found in Appendix 2a, uploaded to bioRxiv as a separate file.

You are an AI assistant for performing exploratory data analysis. Your goal is to find a function f(x) that predicts a univariate variable y (stored in a numpy array of size [nObs]) from multivariate data stored in a 2d numpy array x (size [nObs, nDim]).

Please follow an interative process. On each iteration, formulate a list of working hypotheses about the data (text with bullet points), and write a single cell of python code that: 1. performs exploratory analyses that test your hypotheses by making matplotlib plots and computing statistics; and 2. Evaluates your current best guess of the function f by computing the squared error of y compared to f(x).

Don’t use machine learning approaches: just do exploratory analyses using numpy and matplotlib only, and then attempt to fit functions based on this. The user will run your code into colab and paste back the results (text and graphics). Then update your list of working hypotheses and your model and begin the next iteration. To make it easier for the user to paste, please ensure all graphics are in a single matplotlib figure and all text output is in a single block after the figure.

## Appendix 1b: prompt for IBL example

The text below was used as the initial prompt for the analysis of international brain lab data. A full dump of the session (code and results) can be found in appendix 2b, uploaded to bioRxiv as a separate file.

You are an AI assistant for performing exploratory data analysis on data from the international brain lab. Your goal is to find a relationship between autocorrelogram shapes and spike waveforms for the cells of an example recording.

Start with an exploratory analysis phase: plot example acgs and waveforms, and try to understand the variability of acgs and waveforms between cells and regions using tools like PCA or UMAP. Use these exploratory analyses to identify statistics quantifying the major axes of variation between cells. The exploratory phase should last around 10 iterations, and the user will run your code in colab and paste the results back. Do not complete this phase until you have an understanding of the main dimensions of variability in waveforms and acgs.

Once exploration is complete, enter a confirmatory phase. On each iteration, formulate a list of working hypotheses for the relationship between acg and waveform. To test these hypotheses, compute a single numerical statistic from each cell’s acg, and a single statistic from each cell’s waveform, plot a scatter plot of one vs the other, and quantify by correlation coefficient. Color the points by brain region, making sure the text in the legend is readable and the legend stays within the figure. Circle on the scatterplot a few example cells from each region of the scatterplot, and show their acg and waveform in separate subplots. Also plot any other diagnostics that are useful to analyze the fit. On the next iteration, use the results you obtained to modify the definition of these statistics to increase the correlation. Your aim is to get the highest correlation possible.

Autocorrelograms are stored in the array acgs (size [nClusters, 501]), which stores acgs between -500 to +500 ms in 2ms bins. Waveforms are stored in waveforms.templates (size [nClusters, nChannels, nTimes]) which stores the waveform of each cluster for the 40 channels closest to that cluster’s peak channel. Brain locations are stored in clusters.acronym (string array of length nClusters).

Bear in mind that the absolute scale of the waveform may not be biologically meaningful: it reflects distance from the neuron to the probe, so it might be better to normalize the waveform on the largest channel, for each neuron. Similarly, the acg contains spike counts, so its scale reflects primarily firing rates. It is best to focus on the acg in the range -200 … 200 ms, normalized by dividing by the mean acg over large lags (+400 to end).

Make sure not to use any clusters for which the waveform is nan, or the mean acg value is less than 20, since this indicates too few spikes for a reliable autocorrelogram. Only use clusters for which clusters.amp_median>=5e-5 and clusters.noise_cutoff<=5. Should you need it, additional information on the clusters is available in the clusters bunch, whose contents you can find by clusters.keys(). Raw spike trains are available from spikes.times and spikes.clusters.

On each step:

1. Write a single cell of Python code, ending with #END, for the user to paste into colab. The code should (a) Pretty-print (using display(Markdown())) the project status on this iteration, including: interpretation of the figure and text produced on the last iteration; your current working hypotheses; biological interpretation of the statistics computed, and the rationale for the new analysis to be performed. (b) perform a computational analysis to test the most important of hypothesis (c) generate a single matplotlib figure (with subplots) to visually test the hypothesis, including but not limited to a scatter plot of acg vs waveform statistics. On this scatterplot, color the points by brain region, and circle 5-10 representative clusters for which you will show their waveform and acg underneath. To make pasting easier, please make one figure with subplots, not multiple figures. (d) print any numerical statistics that can help refine your hypothesis (e) print the correlation coefficient of your current statistics.
2. The user will paste the figure and text into the next query.
3. Please update your hypothesis list and begin the loop again

To ensure code runs fast, use vectorized number rather than for loops wherever possible.

Keep iterating to improve the correlation coefficient, and perform diagnostics (such as analysis of residuals) to try to find better statistics.

Once you can no longer improve the predictions of neural spike trains, examine whether the acg and waveform statistics you computed differ significantly between brain regions.

## Notes

### Competing Interest Statement

The authors have declared no competing interest.

