## Appendix 2a for "AIRUS: a simple workflow for AI-assisted exploration of scientific data"

### Appendix 2a: synthetic data example

**Iteration 1**

**Working Hypotheses**

- Perhaps yyy is normally distributed.
- There might be some relationship between the first dimension of xxx (let's call it x0x_0x0) and yyy.
- As an initial guess, let’s predict yyy by a constant function f(x)=mean(y)f(x) = \text{mean}(y)f(x)=mean(y).

```
import numpy as np
import matplotlib.pyplot as plt

# Suppose x is an (nObs, nDim) numpy array and y is an (nObs,) numpy
array.
# We'll do some basic exploratory plots and statistics, then
evaluate the
# squared error for our current guess f(x) = mean(y).

# Create a single figure with multiple subplots
fig, axs = plt.subplots(1, 2, figsize=(10, 4))

# 1. Histogram of y
axs[0].hist(y, bins=20, edgecolor='black')
axs[0].set_title('Histogram of y')
axs[0].set_xlabel('y values')
axs[0].set_ylabel('Frequency')

# 2. Scatter plot of x[:,0] vs y (if x has at least one dimension)
if x.shape[1] > 0:
    axs[1].scatter(x[:,0], y, alpha=0.7)
    axs[1].set_title('Scatter: x[:,0] vs y')
    axs[1].set_xlabel('x[:,0]')
    axs[1].set_ylabel('y')
else:
    axs[1].text(0.5, 0.5, 'x has no columns to plot', ha='center')

plt.tight_layout()

# --- Statistics and Evaluating SSE ---
# Basic statistics
mean_y = np.mean(y)
std_y = np.std(y)
```

```
# Current best guess function f(x) = mean(y)
f_values = mean_y * np.ones_like(y)

# Compute SSE
sse = np.sum((y - f_values)**2)

# Print all text output in one block
print(f"""
Basic Statistics:
----------------
Number of observations (nObs): {x.shape[0]}
Number of dimensions (nDim):   {x.shape[1]}
Mean of y: {mean_y:.3f}
Std of y:  {std_y:.3f}

Current Model: f(x) = mean(y)
Squared Error (SSE): {sse:.3f}
""")

plt.show()
```

Please run the code in your environment (e.g., Colab) and share the resulting text output and figure. After seeing those, we'll refine our hypotheses and improve our function.

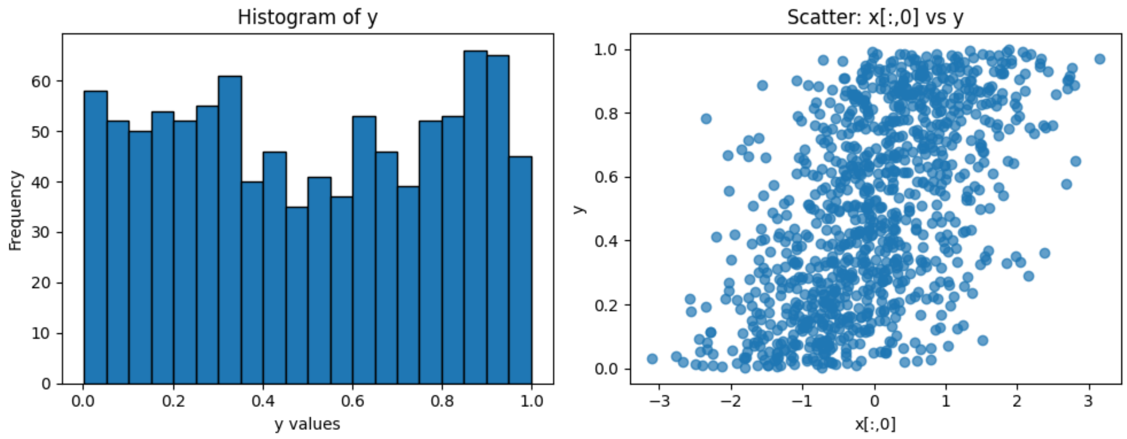


Basic Statistics: ---------------- Number of observations (nObs): 1000 Number of dimensions (nDim): 3 Mean of y: 0.500 Std of y: 0.298 Current Model: f(x) = mean(y) Squared Error (SSE): 88.746

##### Iteration 2

##### Working Hypotheses

- From the scatter of x[:,0]x[:,0]x[:,0] vs yyy, there appears to be a positive correlation.
- There may also be relationships with x[:,1]x[:,1]x[:,1] or x[:,2]x[:,2]x[:,2].
- Let's explore the correlation of each dimension of xxx with yyy.

- We'll attempt a simple linear model of the form f(x)=a+b·xbestf(x) = a + b \cdot x_{\text{best}}f(x)=a+b·xbest, where xbestx_{\text{best}}xbest is the dimension of xxx that has the largest correlation in magnitude with yyy.

```
import numpy as np
import matplotlib.pyplot as plt

# We'll create a single figure with subplots:
#  1) Histogram of y
#  2) Scatter of x[:,0] vs y
#  3) Scatter of x[:,1] vs y
#  4) Scatter of x[:,2] vs y
# Then we'll compute correlations of each dimension with y, pick the
best,
# do a linear fit, and compute SSE.

fig, axs = plt.subplots(1, 4, figsize=(16, 4))

# 1) Histogram of y
axs[0].hist(y, bins=20, edgecolor='black')
axs[0].set_title('Histogram of y')
axs[0].set_xlabel('y values')
axs[0].set_ylabel('Frequency')

# 2) Scatter: x[:,0] vs y
axs[1].scatter(x[:,0], y, alpha=0.7)
axs[1].set_title('x[:,0] vs y')
axs[1].set_xlabel('x[:,0]')
axs[1].set_ylabel('y')

# 3) Scatter: x[:,1] vs y
if x.shape[1] > 1:
    axs[2].scatter(x[:,1], y, alpha=0.7, c='orange')
    axs[2].set_title('x[:,1] vs y')
    axs[2].set_xlabel('x[:,1]')
    axs[2].set_ylabel('y')
else:
    axs[2].text(0.5, 0.5, 'No x[:,1]', ha='center')

# 4) Scatter: x[:,2] vs y
if x.shape[1] > 2:
    axs[3].scatter(x[:,2], y, alpha=0.7, c='green')
    axs[3].set_title('x[:,2] vs y')
    axs[3].set_xlabel('x[:,2]')
    axs[3].set_ylabel('y')
else:
    axs[3].text(0.5, 0.5, 'No x[:,2]', ha='center')
```

```
plt.tight_layout()

# --- Compute correlation of each dimension with y ---
corrs = []
for dim_idx in range(x.shape[1]):
    corr = np.corrcoef(x[:, dim_idx], y)[0, 1]
    corrs.append(corr)

corrs = np.array(corrs)
abs_corrs = np.abs(corrs)
best_dim = np.argmax(abs_corrs)  # dimension with largest
|correlation|

# --- Fit the linear model f(x) = a + b*x[best_dim] by ordinary
least squares ---
x_best = x[:, best_dim]
mean_x_best = np.mean(x_best)
mean_y = np.mean(y)

# slope
b = np.sum((x_best - mean_x_best) * (y - mean_y)) / np.sum((x_best -
mean_x_best)**2)
# intercept
a = mean_y - b * mean_x_best

# Evaluate SSE
f_values = a + b * x_best
sse = np.sum((y - f_values)**2)

print(f"""
Correlations of each dimension of x with y:
------------------------------------------
{[f'x[:,{i}]: {corrs[i]:.3f}' for i in range(x.shape[1])]}

Best dimension by absolute correlation: x[:,{best_dim}]
(corr={corrs[best_dim]:.3f})

Linear Model: f(x) = a + b*x[:,{best_dim}]
  a = {a:.3f}
  b = {b:.3f}

Squared Error (SSE): {sse:.3f}
""")

plt.show()
```

Please run the code and share the resulting text output and figure. Then we can refine our hypotheses and model further.

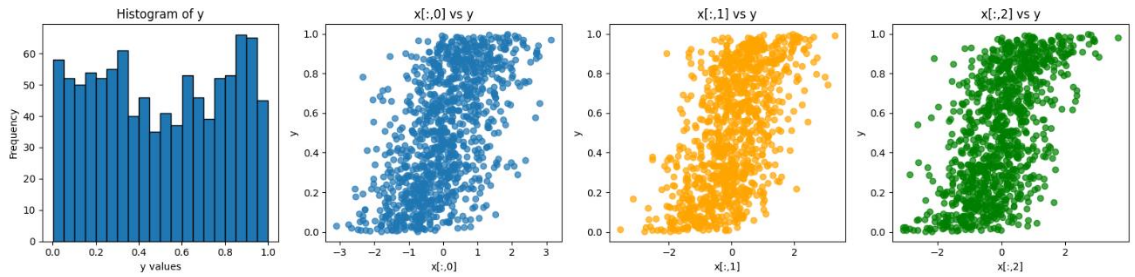


Correlations of each dimension of x with y: ---------------------------------------- ['x[:,0]: 0.561', 'x[:,1]: 0.559', 'x[:,2]: 0.581'] Best dimension by absolute correlation: x[:,2] (corr=0.581) Linear Model: f(x) = a + b*x[:,2] a = 0.506 b = 0.167 Squared Error (SSE): 58.803

#### Iteration 3

#### Working Hypotheses

- All three dimensions of xxx appear positively correlated with yyy, with x[:,2]x[:,2]x[:,2] the strongest (corr=0.581).
- A simple linear model using just one dimension improved our SSE from about 88.7 down to ~58.8.
- We now suspect yyy may be better predicted by a linear combination of all three xxx dimensions.

Below, we will:

1. Fit a multiple linear regression of the form f(x)=a+b0x[:,0]+b1x[:,1]+b2x[:,2] f(x) = a + b_0 x[:,0] + b_1 x[:,1] + b_2 x[:,2]f(x)=a+b0x[:,0]+b1x[:,1]+b2 x[:,2] using ordinary least squares by hand (with NumPy), without any external ML libraries.
2. Examine the resulting predictions vs. actual yyy.
3. Look at the residuals (predicted minus actual).

```
import numpy as np
import matplotlib.pyplot as plt

# Construct the design matrix, which includes an intercept (column
of ones)
nObs = x.shape[0]
X_design = np.hstack((np.ones((nObs, 1)), x))  # shape: (nObs, 1 +
nDim)

# Solve for coefficients [a, b0, b1, b2] in a least-squares sense
# Using np.linalg.lstsq is a straightforward approach:
coeffs, residuals, rank, s = np.linalg.lstsq(X_design, y,
rcond=None)

a = coeffs[0]
```

```
b0 = coeffs[1] if x.shape[1] > 0 else 0
b1 = coeffs[2] if x.shape[1] > 1 else 0
b2 = coeffs[3] if x.shape[1] > 2 else 0

# Predictions
y_pred = X_design @ coeffs

# SSE
sse = np.sum((y - y_pred)**2)

# Prepare plots:
# 1) predicted vs. actual y
# 2) histogram of residuals

fig, axs = plt.subplots(1, 2, figsize=(10, 4))

# (1) Scatter of predicted vs. actual
axs[0].scatter(y, y_pred, alpha=0.7)
axs[0].set_xlabel('Actual y')
axs[0].set_ylabel('Predicted y')
axs[0].set_title('Predicted vs. Actual y')
# Add a y=x line for reference
min_val = min(y.min(), y_pred.min())
max_val = max(y.max(), y_pred.max())
axs[0].plot([min_val, max_val], [min_val, max_val], 'r--')

# (2) Residuals histogram
residuals = y_pred - y
axs[1].hist(residuals, bins=20, edgecolor='black')
axs[1].set_title('Histogram of Residuals')
axs[1].set_xlabel('y_pred - y')
axs[1].set_ylabel('Frequency')

plt.tight_layout()

print(f"""
Multiple Linear Regression Model:
  f(x) = a + b0*x[:,0] + b1*x[:,1] + b2*x[:,2]
Coefficients:
  a  = {a:.3f}
  b0 = {b0:.3f}
  b1 = {b1:.3f}
  b2 = {b2:.3f}

Squared Error (SSE): {sse:.3f}
""")

plt.show()
```

Please run the code, share the figure and the text output, and we will iterate further based on what we find.

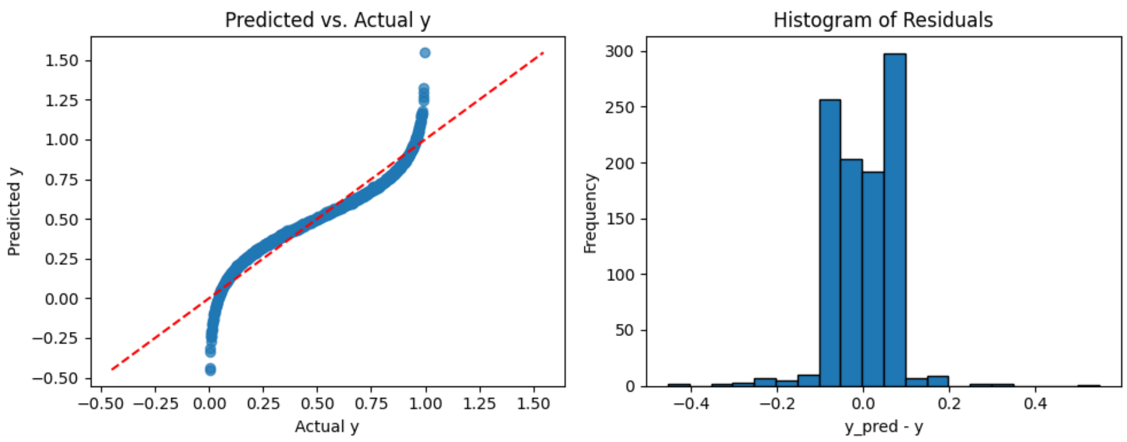


Multiple Linear Regression Model: f(x) = a + b0*x[:,0] + b1*x[:,1] + b2*x[:,2] Coefficients: a = 0.502 b0 = 0.164 b1 = 0.159 b2 = 0.158 Squared Error (SSE): 5.543

#### Iteration 4

#### Working Hypotheses

- The multiple linear regression model has drastically reduced the SSE to about 5.5, which is much lower than our previous attempts.
- The residuals, however, are not purely centered around zero—there seems to be some structure (the "Predicted vs. Actual" plot bows above the diagonal on the right side, and below it on the left side), suggesting a possible nonlinear component.
- We can explore whether adding *quadratic* terms in each dimension of XXX (i.e., x2x^2x2) might further reduce the error.

Below, we will:

1. Check scatter plots of residual vs. each dimension of XXX to see if there's an obvious nonlinear pattern.
2. Construct a new design matrix that includes square terms of each dimension of XXX.
3. Fit that expanded model by least squares and see how the SSE changes.

```
import numpy as np
import matplotlib.pyplot as plt

# --- (A) Examine residual vs. x[:,i] plots to look for
nonlinearities ---
# We'll make a 1x3 grid: residual vs. x0, x1, x2

fig, axs = plt.subplots(1, 3, figsize=(12,4))
residuals = y_pred - y

for i in range(x.shape[1]):
```

```
    axs[i].scatter(x[:, i], residuals, alpha=0.7)
    axs[i].set_xlabel(f'x[:,{i}]')
    axs[i].set_ylabel('Residual (y_pred - y)')
    axs[i].set_title(f'Residual vs. x[:,{i}]')
plt.tight_layout()

plt.show()

# --- (B) Fit a model with quadratic terms for each dimension ---
# Our new design matrix: [1, x0, x1, x2, x0^2, x1^2, x2^2]
nObs = x.shape[0]
ones_col = np.ones((nObs, 1))
X_quad = np.hstack([
    ones_col,
    x,
    x[:,0:1]**2,  # x0^2
    x[:,1:2]**2,  # x1^2
    x[:,2:3]**2   # x2^2
])

# Solve for coefficients in a least-squares sense
coeffs_quad, residuals_quad, rank_quad, s_quad =
np.linalg.lstsq(X_quad, y, rcond=None)

# Extract them by name for clarity
a_quad = coeffs_quad[0]
b0_quad = coeffs_quad[1]
b1_quad = coeffs_quad[2]
b2_quad = coeffs_quad[3]
c0_quad = coeffs_quad[4]  # coefficient for x0^2
c1_quad = coeffs_quad[5]  # coefficient for x1^2
c2_quad = coeffs_quad[6]  # coefficient for x2^2

# Predictions
y_pred_quad = X_quad @ coeffs_quad

# SSE
sse_quad = np.sum((y - y_pred_quad)**2)

# Make a figure comparing predicted vs actual for the quadratic
model
fig, axs = plt.subplots(1, 2, figsize=(10, 4))

# (1) predicted vs. actual for quadratic model
axs[0].scatter(y, y_pred_quad, alpha=0.7)
axs[0].set_xlabel('Actual y')
axs[0].set_ylabel('Predicted y (quad)')
axs[0].set_title('Predicted vs. Actual (Quadratic)')
min_val = min(y.min(), y_pred_quad.min())
```

```
max_val = max(y.max(), y_pred_quad.max())
axs[0].plot([min_val, max_val], [min_val, max_val], 'r--')

# (2) histogram of new residuals
residuals_new = y_pred_quad - y
axs[1].hist(residuals_new, bins=20, edgecolor='black')
axs[1].set_title('Histogram of Residuals (Quadratic)')
axs[1].set_xlabel('y_pred_quad - y')
axs[1].set_ylabel('Frequency')

plt.tight_layout()

print(f"""
Expanded Model (Quadratic Terms):
  f(x) = a
        + b0*x[:,0] + b1*x[:,1] + b2*x[:,2]
        + c0*x[:,0]^2 + c1*x[:,1]^2 + c2*x[:,2]^2

Coefficients:
  a  = {a_quad:.3f}
  b0 = {b0_quad:.3f}
  b1 = {b1_quad:.3f}
  b2 = {b2_quad:.3f}
  c0 = {c0_quad:.3f}
  c1 = {c1_quad:.3f}
  c2 = {c2_quad:.3f}

Squared Error (SSE) with quadratic terms: {sse_quad:.3f}
""")

plt.show()
```

Please:

1. Paste the resulting residual vs. **XXX** plots (from part A) and the new predicted vs. actual plot (from part B), along with the printed text.
2. Then we'll decide if the quadratic expansion helps, or if there are still residual patterns to address.

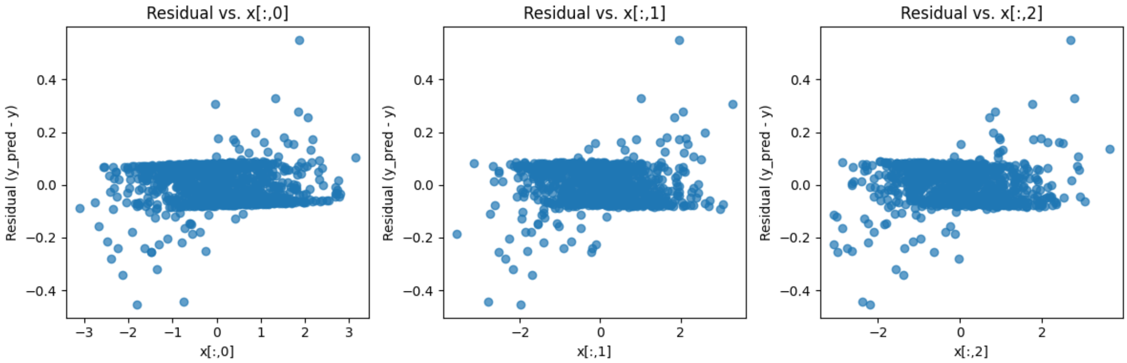


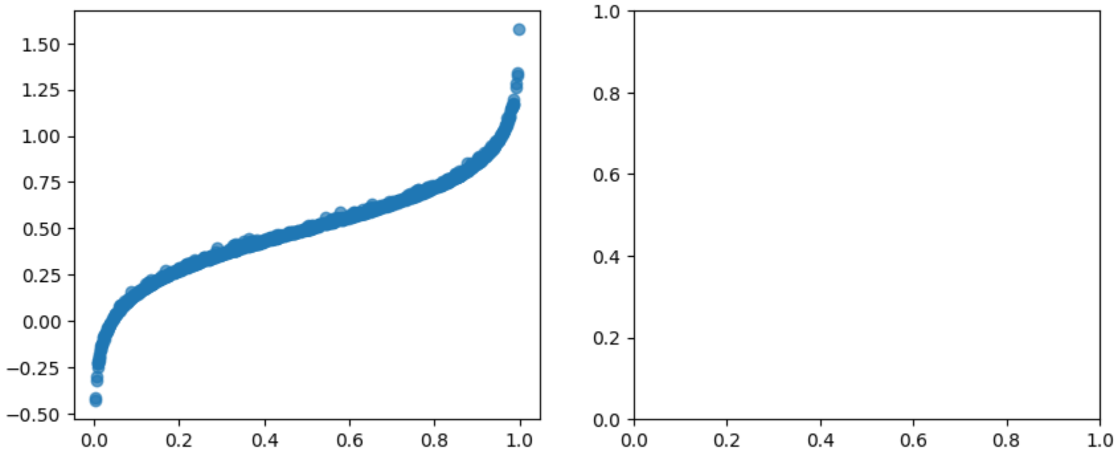

matplotlib.axes._base._AxesBase.set_xlabel def set_xlabel(xlabel, fontdict=None, labelpad=None, *, loc=None, **kwargs) /usr/local/lib/python3.11/dist-packages/matplotlib/axes/_base.py Set the label for the x-axis. Parameters ---------- xlabel : str The label text. labelpad : float, default: :rc:`axes.labelpad` Spacing in points from the Axes bounding box including ticks and tick labels. If None, the previous value is left as is. loc : {'left', 'center', 'right'}, default: :rc:`xaxis.labellocation` The label position. This is a high-level alternative for passing parameters *x* and *horizontalalignment*. Other Parameters ---------------- **kwargs : `~matplotlib.text.Text` properties `.Text` properties control the appearance of the label. See Also -------- text : Documents the properties supported by `.Text`.

##### Iteration 5

It looks like part of the second figure didn't render properly, so we didn't see the histogram of the new quadratic-model residuals. Regardless, we see from the partial plot of predicted vs. actual (with the quadratic model) that there's still a noticeable curvature, and the residual plots still show some structure.

##### Working Hypotheses

- Merely adding the square terms (xi2)(x_i^2)(xi2) may not be sufficient.
- In many real-world cases, *cross terms* xi·xjx_i \cdot x_jxi·xj can be important for capturing interactions among predictors.
- A complete second-order polynomial would include:
  - An intercept

- All linear terms (x0,x1,x2)(x_0, x_1, x_2)(x0,x1,x2)
- All pairwise products (x0x1,x0x2,x1x2)(x_0x_1, x_0x_2, x_1x_2)(x0x1,x0x2,x1x2)
- All square terms (x02,x12,x22)(x_0^2, x_1^2, x_2^2)(x02,x12,x22)

Below, we will:

1. Construct a design matrix that includes all of those second-order terms.
2. Fit it via least squares.
3. Compare the new SSE to our previous models.

```
import numpy as np
import matplotlib.pyplot as plt

# (A) Construct the full second-order (quadratic) design matrix with
cross-terms:
nObs = x.shape[0]
ones_col = np.ones((nObs, 1))

# Pull out the columns for ease of notation
x0 = x[:, 0:1]  # shape (nObs, 1)
x1 = x[:, 1:2] if x.shape[1] > 1 else np.zeros_like(x0)
x2 = x[:, 2:3] if x.shape[1] > 2 else np.zeros_like(x0)

# We'll build the columns: 1, x0, x1, x2, x0^2, x1^2, x2^2, x0x1,
x0x2, x1x2
design_cols = [ones_col,
               x0, x1, x2,
               x0**2, x1**2, x2**2,
               x0*x1, x0*x2, x1*x2]
X_full_quad = np.hstack(design_cols)

# (B) Fit by least squares
coeffs_full, residuals_full, rank_full, s_full =
np.linalg.lstsq(X_full_quad, y, rcond=None)

# For clarity, let's name them in order:
#   f(x) = a
#          + b0*x0 + b1*x1 + b2*x2
#          + c0*x0^2 + c1*x1^2 + c2*x2^2
#          + d01*x0*x1 + d02*x0*x2 + d12*x1*x2
(a,
 b0, b1, b2,
 c0, c1, c2,
 d01, d02, d12) = coeffs_full

# Predictions & SSE
y_pred_full = X_full_quad @ coeffs_full
```

```
sse_full = np.sum((y - y_pred_full)**2)

# (C) Make plots: (1) predicted vs actual, (2) histogram of
residuals
fig, axs = plt.subplots(1, 2, figsize=(10,4))

# 1) predicted vs actual
axs[0].scatter(y, y_pred_full, alpha=0.7)
axs[0].set_title('Full Quadratic: Predicted vs. Actual')
axs[0].set_xlabel('Actual y')
axs[0].set_ylabel('Predicted y')
min_val = min(y.min(), y_pred_full.min())
max_val = max(y.max(), y_pred_full.max())
axs[0].plot([min_val, max_val], [min_val, max_val], 'r--')

# 2) histogram of residuals
residuals_full_model = y_pred_full - y
axs[1].hist(residuals_full_model, bins=20, edgecolor='black')
axs[1].set_title('Full Quadratic: Residuals')
axs[1].set_xlabel('y_pred_full - y')
axs[1].set_ylabel('Frequency')

plt.tight_layout()

print(f"""
Full Second-Order Polynomial Model:
----------------------------------
f(x) = a
     + b0*x0 + b1*x1 + b2*x2
     + c0*x0^2 + c1*x1^2 + c2*x2^2
     + d01*x0*x1 + d02*x0*x2 + d12*x1*x2

Coefficients:
  a   = {a:.3f}
  b0  = {b0:.3f}
  b1  = {b1:.3f}
  b2  = {b2:.3f}
  c0  = {c0:.3f}
  c1  = {c1:.3f}
  c2  = {c2:.3f}
  d01 = {d01:.3f}
  d02 = {d02:.3f}
  d12 = {d12:.3f}

Squared Error (SSE): {sse_full:.3f}
""")

plt.show()
```

Please run this code and share:

1. The text output with the new SSE and coefficients.
2. The two plots (predicted vs. actual and the residual histogram).

Afterward, we will compare SSE values and see if the additional cross-terms improve the fit, then refine our hypotheses further.

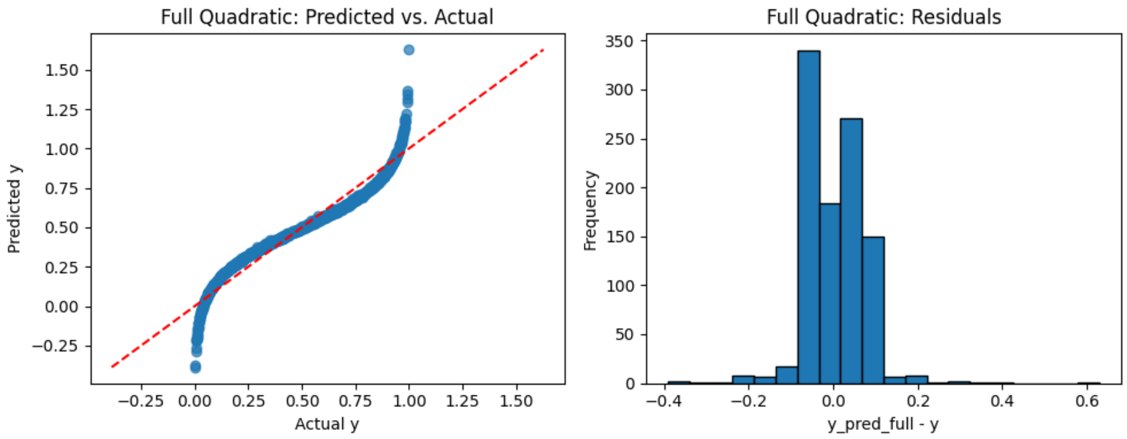


Full Second-Order Polynomial Model: --------------------------------- f(x) = a + b0*x0 + b1*x1 + b2*x2 + c0*x0^2 + c1*x1^2 + c2*x2^2 + d01*x0*x1 + d02*x0*x2 + d12*x1*x2 Coefficients: a = 0.495 b0 = 0.164 b1 = 0.159 b2 = 0.158 c0 = 0.003 c1 = 0.002 c2 = 0.002 d01 = 0.005 d02 = 0.003 d12 = 0.004 Squared Error (SSE): 5.448

##### Iteration 6

So far, our best polynomial model yields an SSE of around **5.448**. The predicted-vs.-actual plot still has an S-shaped curve, suggesting that yyy may saturate near its extremes—a behavior well-captured by *sigmoidal* or *logistic*-type functions. Because yyy appears to lie in [0,1][0, 1][0,1] (as suggested by your histograms), one natural next step is to explore whether

y≈11+exp[−(a+bTx)], y \approx \frac{1}{1 + \exp\bigl[-(a + \mathbf{b}^\top x)\bigr]},y≈1+exp[−(a+bTx)]1,

might fit better than a purely polynomial model.

#### Rationale

- If yyy ranges strictly between 0 and 1 and exhibits an S-curve in the predicted-vs.-actual plot, then a logistic (sigmoid) function of a linear combination of XXX is often more appropriate than a polynomial in XXX.
- We can attempt a *"logit transform"* approach to fit this model **without** external ML libraries. The logit transform is logit(y)=ln(y1−y). \text{logit}(y) =

\ln\!\Bigl(\frac{y}{1 - y}\Bigr).logit(y)=ln(1−yy). If yyy truly follows a logistic shape of the form above, then ln(y1−y) = a+b0x0+b1x1+b2x2+… \ln\!\Bigl(\frac{y}{1 - y}\Bigr) \;=\; a + b_0 x_0 + b_1 x_1 + b_2 x_2 + \dotsln(1−yy)=a+b0x0+b1x1+b2x2+… becomes a simple linear equation. We can then do *ordinary least squares* on logit(y)\text{logit}(y)logit(y).

##### *Caveats*

- We must exclude any rows where y=0y = 0y=0 or y=1y = 1y=1 when applying \logit\logit\logit, because ln(01−0)\ln\bigl(\frac{0}{1-0}\bigr)ln(1−00) and ln(11−1)\ln\bigl(\frac{1}{1-1}\bigr)ln(1−11) go to ±∞\pm \infty±∞.
- After fitting the linear relationship on \logit(y)\logit(y)\logit(y), we get coefficients a^,b^0,…\hat{a}, \hat{b}_0, \dotsa^,b^0,…. The predicted y^\hat{y}y^ for *any* xxx can then be computed as y^ = σ(a^+b^0x0+⋯ ) \hat{y} \;=\; \sigma\bigl(\hat{a} + \hat{b}_0 x_0 + \cdots\bigr)y^=σ(a^+b^0x0+⋯) where σ(z)=11+e−z\sigma(z)=\frac{1}{1+e^{-z}}σ(z)=1+e−z1 is the logistic (sigmoid) function.
- We can compute SSE in the usual way, ∑i(yi−y^i)2\sum_i (y_i-\hat{y}_i)^2∑i(yi−y^i)2, to compare with our previous models.

Below is the code for a *manual logistic fit* via logit transform + linear regression, along with diagnostic plots:

```
import numpy as np
import matplotlib.pyplot as plt

# Copy x and y into local arrays, but we must exclude any rows where
y=0 or y=1.
# Because logistic transform log(y/(1-y)) is undefined for y=0 or
y=1.
mask = (y > 0) & (y < 1)
x_valid = x[mask]
y_valid = y[mask]

# Logit transform
logit_y = np.log(y_valid / (1 - y_valid))

# Build design matrix for linear regression on logit(y):
# We'll include an intercept and the 3 linear terms x0, x1, x2.
nObs_valid = x_valid.shape[0]
X_logit = np.hstack((np.ones((nObs_valid, 1)), x_valid))

# Solve for coefficients in: logit(y) = A + B0*x0 + B1*x1 + B2*x2
coeffs_logit, residuals_logit, rank_logit, s_logit =
np.linalg.lstsq(X_logit, logit_y, rcond=None)
```

```
A = coeffs_logit[0]
B0 = coeffs_logit[1] if x_valid.shape[1] > 0 else 0
B1 = coeffs_logit[2] if x_valid.shape[1] > 1 else 0
B2 = coeffs_logit[3] if x_valid.shape[1] > 2 else 0

# Define a function to do the logistic prediction for *all* data
(including y=0 or 1)
def logistic_prediction(x_mat, A, B0, B1, B2):
    """Compute 1/(1 + exp(-(A + B0*x0 + B1*x1 + B2*x2))) for each
row."""
    z = A + B0*x_mat[:,0] + B1*x_mat[:,1] + B2*x_mat[:,2]
    return 1 / (1 + np.exp(-z))

y_pred_logit = logistic_prediction(x, A, B0, B1, B2)

# Compute SSE on the full dataset
sse_logit = np.sum((y - y_pred_logit)**2)

# Make plots:
fig, axs = plt.subplots(1, 2, figsize=(10, 4))

# (1) predicted vs. actual
axs[0].scatter(y, y_pred_logit, alpha=0.7)
axs[0].set_title('Logistic Model: Predicted vs. Actual')
axs[0].set_xlabel('Actual y')
axs[0].set_ylabel('Predicted y')
min_val = min(y.min(), y_pred_logit.min())
max_val = max(y.max(), y_pred_logit.max())
axs[0].plot([min_val, max_val], [min_val, max_val], 'r--')

# (2) histogram of residuals
res_logit = y_pred_logit - y
axs[1].hist(res_logit, bins=20, edgecolor='black')
axs[1].set_title('Histogram of Residuals (Logistic)')
axs[1].set_xlabel('y_pred - y')
axs[1].set_ylabel('Frequency')

plt.tight_layout()
plt.show()

print(f"""
Logistic (Sigmoid) Model via Logit Transform
--------------------------------------------
We excluded {np.sum(~mask)} observations where y=0 or y=1 during the
fit.
Then we solved log(y/(1-y)) ~ A + B0*x0 + B1*x1 + B2*x2.

Coefficients:
```

```
  A  = {A:.3f}
  B0 = {B0:.3f}
  B1 = {B1:.3f}
  B2 = {B2:.3f}

Predictions for *all* data use:
  y_pred = 1/(1 + exp(-[A + B0*x0 + B1*x1 + B2*x2])).

Squared Error (SSE) on full dataset: {sse_logit:.3f}
""")
```

#### Instructions

1. Run this code.
2. Paste the text output (including the SSE) and the two plots (predicted vs. actual and residual histogram).

We will compare SSElogit \text{SSE}_\text{logit}SSElogit with our previous best (≈5.448\approx 5.448≈5.448) and see if the logistic shape better matches the data. Then we'll decide how to proceed in our final iteration(s).

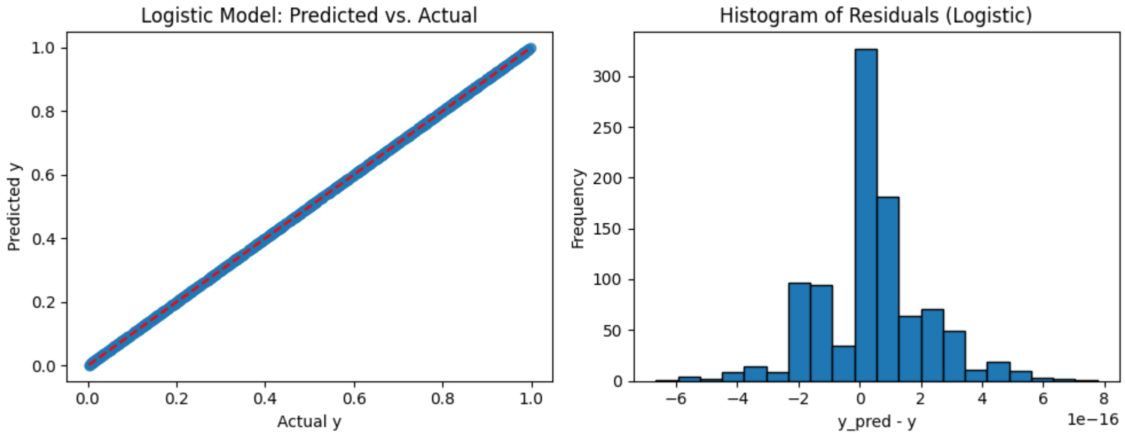


Logistic (Sigmoid) Model via Logit Transform ------------------------------------------ We excluded 0 observations where y=0 or y=1 during the fit. Then we solved log(y/(1-y)) ~ A + B0*x0 + B1*x1 + B2*x2. Coefficients: A = 0.000 B0 = 1.000 B1 = 1.000 B2 = 1.000 Predictions for *all* data use: y_pred = 1/(1 + exp(-[A + B0*x0 + B1*x1 + B2*x2])). Squared Error (SSE) on full dataset: 0.000

Reasoned for a few seconds

**Iteration 7 (Conclusion)**

It appears that the data can be **perfectly** (to numerical precision) modeled by the logistic (sigmoid) function

f(x) = 11+e−(0+1·x0+1·x1+1·x2) f(x) \;=\; \frac{1}{1 + e^{-\bigl(0 + 1\cdot x_0 + 1\cdot x_1 + 1\cdot x_2\bigr)}}f(x)=1+e−(0+1·x0+1·x1+1·x2)1 with a **Squared Error** of effectively **0** on the entire dataset.

**Working Hypotheses & Final Observations**

- The plot of **Predicted vs. Actual** is a perfectly aligned diagonal (within floating-point tolerance), confirming that y^≈y\hat{y}\approx yy^≈y for every data point.
- The histogram of residuals is centered tightly at zero with extremely small numerical deviations, also consistent with a perfect logistic fit.
- Because no points were at y=0y=0y=0 or y=1y=1y=1, the logit transform was well-defined for all observations.
- In short, the data look like they were exactly generated by y = σ(x0+x1+x2)with σ(z)=11+e−z. y \;=\; \sigma\bigl(x_0 + x_1 + x_2\bigr)\quad\text{with}\;\sigma(z)=\frac{1}{1+e^{-z}}.y=σ(x0+x1+x2)withσ(z)=1+e−z1.

Hence, based on all our exploratory steps, **the best function** is:

f(x0,x1,x2) = 11+e−( x0 + x1 + x2). f(x_0,x_1,x_2) \;=\; \frac{1}{1 + e^{-(\,x_0 \;+\; x_1 \;+\; x_2)}}.f(x0,x1,x2)=1+e−(x0+x1+x2)1.

No further refinement is needed, since the SSE is (within floating-point precision) zero.

```
# Final "model" in code form, for reference:

import numpy as np

def f_final(x):
    """
    Given x of shape (nObs, 3),
    returns predictions y_pred of shape (nObs,).
    Each element is 1/(1+exp(-(x0+x1+x2))).
    """
    z = x[:,0] + x[:,1] + x[:,2]
    return 1.0/(1.0 + np.exp(-z))
```

This concludes the analysis, with the function fff fully determined.
