## Appendix 2b for "AIRUS: a simple workflow for AI-assisted exploration of scientific data"

### Appendix 2b: relating autocorrelograms to spike trains

#### Code run to load data and compute acgs (not shown to ai)

This code was run to load the data from an example recording, prior to beginning the workflow. The AI did not have access to the code.

In [15]:

```
!pip install ibllib
!pip install umap-learn
```

In [16]:

```
from one.api import ONE
ONE.setup(base_url='https://openalyx.internationalbrainlab.org',
silent=True)
one = ONE(password='international')
from brainbox.singlecell import acorr
from one.api import ONE
from brainbox.io.one import SpikeSortingLoader
from iblatlas.regions import BrainRegions
from iblatlas.atlas import AllenAtlas
ba = AllenAtlas()

pid = '1a276285-8b0e-4cc9-9f0a-a3a002978724'
ssl = SpikeSortingLoader(pid=pid, one=one, atlas=ba)
spikes, clusters, channels = ssl.load_spike_sorting(revision='2024-
05-06')
clusters = ssl.merge_clusters(spikes, clusters, channels)
waveforms = ssl.load_spike_sorting_object('waveforms')  # loads in
the template waveforms
nClusters = len(clusters.acronym)
```

**Connected to https://openalyx.internationalbrainlab.org as user "intbrainlab"**

In [17]:

```
import numpy as np
NBINS = 501 # including center bin
WIDTH = 1 # second total
acgs = np.zeros((nClusters, NBINS))
print(f'{nClusters} to compute for pid {pid}')
for i in range(nClusters):
  acgs[i,:] = acorr(spikes.times[spikes.clusters==i], WIDTH/(NBINS-
1), WIDTH)
  if (i%100)==0: print('.', end='')
```

**1143 to compute for pid 1a276285-8b0e-4cc9-9f0a-a3a002978724**
**............**

#### Results of Iteration

The text below shows the AI-generated code that was run in colab, and the results it produced. User interventions, which start later in the iterative process are shown in red.

In [18]:

```
# First Exploratory Iteration

import numpy as np
import matplotlib.pyplot as plt
from IPython.display import Markdown, display
from sklearn.decomposition import PCA

# For nicer plotting aesthetics
plt.style.use('default')

#==================================================================
# (a) Display project status
#==================================================================
display(Markdown("""
**Exploratory Analysis - Iteration 1**

**Interpretation of Previous Results**
_(No previous plots yet, this is our first iteration.)_

**Current Working Hypotheses**
1. The shape of each neuron's autocorrelogram (ACG) varies primarily
with its firing pattern (e.g. bursting vs. regular spiking).
2. The shape of each neuron's waveform varies primarily with cell
type (e.g. narrow-spiking vs. broad-spiking).
3. We suspect there could be multiple dimensions of variability in
both ACGs and waveforms, possibly tied to brain region.

**Biological Interpretation**
- ACG shape can reflect firing dynamics (e.g., burstiness).
- Waveform shape often reflects electrophysiological characteristics
of the cell (e.g., fast-spiking interneuron vs excitatory neuron).

**Rationale for New Analysis**
In this first step, we will:
1. Filter out unreliable clusters (insufficient spikes, poor
quality, or NaN waveforms).
2. Normalize the ACG (focusing on -200ms to +200ms, and scaling by
the mean at large lags).
```

3. Collapse each neuron's waveform to a single channel (the channel with largest absolute amplitude) and normalize.
4. Perform PCA separately on the set of normalized waveforms and normalized ACGs to visualize major axes of variation.
5. Check the initial correlation (as a placeholder measure) between the first PC of waveforms and the first PC of ACGs.

```
"""))

#==================================================================
# (b) Perform the computational analysis
#==================================================================

# --- Assume we already have these objects in memory ---
# acgs: shape [nClusters, 501]
# waveforms.templates: shape [nClusters, nChannels, nTimes]
# clusters.acronym: shape [nClusters] (array of region labels)
# clusters.amp_median: shape [nClusters] (median amplitude)
# clusters.noise_cutoff: shape [nClusters] (noise metrics)

# 1) Filter out clusters with bad waveforms or insufficient spikes
#    - Must not have NaN waveforms
#    - Mean of acg >= 20
#    - clusters.amp_median >= 5e-5
#    - clusters.noise_cutoff <= 5

valid_waveform_mask = ~np.isnan(waveforms.templates).any(axis=(1,2))
mean_acg = acgs.mean(axis=1)
valid_acg_mask = mean_acg >= 20
valid_amp_mask = clusters.amp_median >= 5e-5
valid_noise_mask = clusters.noise_cutoff <= 5

valid_mask = valid_waveform_mask & valid_acg_mask & valid_amp_mask &
valid_noise_mask
valid_indices = np.where(valid_mask)[0]

# Subset arrays
acgs_valid = acgs[valid_indices, :]
waveforms_valid = waveforms.templates[valid_indices, :, :]
regions_valid = clusters.acronym[valid_indices]

# 2) Normalize ACG
#    Focus on -200..200 ms => index 150:351 (2 ms per bin,
center=250)
#    Divide by mean at large lags => index 400:501
acg_subset = acgs_valid[:, 150:351]  # shape [nValid, 201]
acg_norm_factor = acgs_valid[:, 400:].mean(axis=1, keepdims=True)  #
shape [nValid, 1]
```

```
acg_norm = acg_subset / (acg_norm_factor + 1e-12)  # to avoid
division by zero

# 3) Collapse waveform to single channel: pick largest amplitude
channel (by peak-to-trough)
#    Then normalize by absolute max. We'll sample across time
dimension [nTimes].
#    waveforms_valid[i, c, :] => waveform of cluster i, channel c,
across time.
peak_ch_idxs = np.argmax(np.max(np.abs(waveforms_valid), axis=2),
axis=1)  # which channel has largest amplitude for each cluster
collapsed_waveforms = []
for i in range(len(valid_indices)):
    w = waveforms_valid[i, peak_ch_idxs[i], :]
    w = w / (np.max(np.abs(w)) + 1e-12)  # normalize
    collapsed_waveforms.append(w)
collapsed_waveforms = np.array(collapsed_waveforms)  # shape
[nValid, nTimes]

# 4) Perform PCA on ACGs and waveforms
#    Flatten each into a single vector to feed PCA.
#    We'll keep 2 PCs for plotting.
pca_acg = PCA(n_components=2)
pca_wave = PCA(n_components=2)

X_acg = acg_norm.reshape(len(acg_norm), -1)  # shape [nValid, 201]
X_wav = collapsed_waveforms.reshape(len(collapsed_waveforms), -1)  #
shape [nValid, nTimes]

acg_scores = pca_acg.fit_transform(X_acg)   # shape [nValid, 2]
wave_scores = pca_wave.fit_transform(X_wav) # shape [nValid, 2]

# 5) Calculate a placeholder correlation: corr between PC1 of
waveforms and PC1 of acg
corr_coefficient = np.corrcoef(acg_scores[:,0],
wave_scores[:,0])[0,1]

#==================================================================
# (c) Generate a single matplotlib figure with subplots
#==================================================================
fig = plt.figure(figsize=(12, 6))

# Subplot 1: PCA of ACG
ax1 = fig.add_subplot(1, 2, 1)
scatter1 = ax1.scatter(acg_scores[:,0], acg_scores[:,1],
c=np.arange(len(acg_scores)), cmap='viridis', s=15)
ax1.set_title('ACG PCA (PC1 vs PC2)')
ax1.set_xlabel('ACG PC1')
ax1.set_ylabel('ACG PC2')
```

```
# Subplot 2: PCA of waveforms
ax2 = fig.add_subplot(1, 2, 2)
scatter2 = ax2.scatter(wave_scores[:,0], wave_scores[:,1],
c=np.arange(len(wave_scores)), cmap='plasma', s=15)
ax2.set_title('Waveform PCA (PC1 vs PC2)')
ax2.set_xlabel('Waveform PC1')
ax2.set_ylabel('Waveform PC2')

plt.tight_layout()
plt.show()

#==================================================================
# (d) Print numeric statistics
#==================================================================
print(f"Number of valid clusters passing all filters:
{len(valid_indices)}")
print(f"Shape of normalized ACG subset: {acg_norm.shape}")
print(f"Shape of collapsed waveforms: {collapsed_waveforms.shape}")
print("Explained variance ratio (ACG PCA):",
pca_acg.explained_variance_ratio_)
print("Explained variance ratio (Waveform PCA):",
pca_wave.explained_variance_ratio_)

#==================================================================
# (e) Print correlation coefficient
#==================================================================
print(f"Correlation between ACG PC1 and Waveform PC1:
{corr_coefficient:.4f}")

#END
```

**Exploratory Analysis - Iteration 1**

**Interpretation of Previous Results**
*(No previous plots yet, this is our first iteration.)*

**Current Working Hypotheses**

1. The shape of each neuron's autocorrelogram (ACG) varies primarily with its firing pattern (e.g. bursting vs. regular spiking).
2. The shape of each neuron's waveform varies primarily with cell type (e.g. narrow-spiking vs. broad-spiking).
3. We suspect there could be multiple dimensions of variability in both ACGs and waveforms, possibly tied to brain region.

**Biological Interpretation**

- ACG shape can reflect firing dynamics (e.g., burstiness).

- Waveform shape often reflects electrophysiological characteristics of the cell (e.g., fast-spiking interneuron vs excitatory neuron).

**Rationale for New Analysis**
In this first step, we will:

1. Filter out unreliable clusters (insufficient spikes, poor quality, or NaN waveforms).
2. Normalize the ACG (focusing on -200ms to +200ms, and scaling by the mean at large lags).
3. Collapse each neuron's waveform to a single channel (the channel with largest absolute amplitude) and normalize.
4. Perform PCA separately on the set of normalized waveforms and normalized ACGs to visualize major axes of variation.
5. Check the initial correlation (as a placeholder measure) between the first PC of waveforms and the first PC of ACGs.

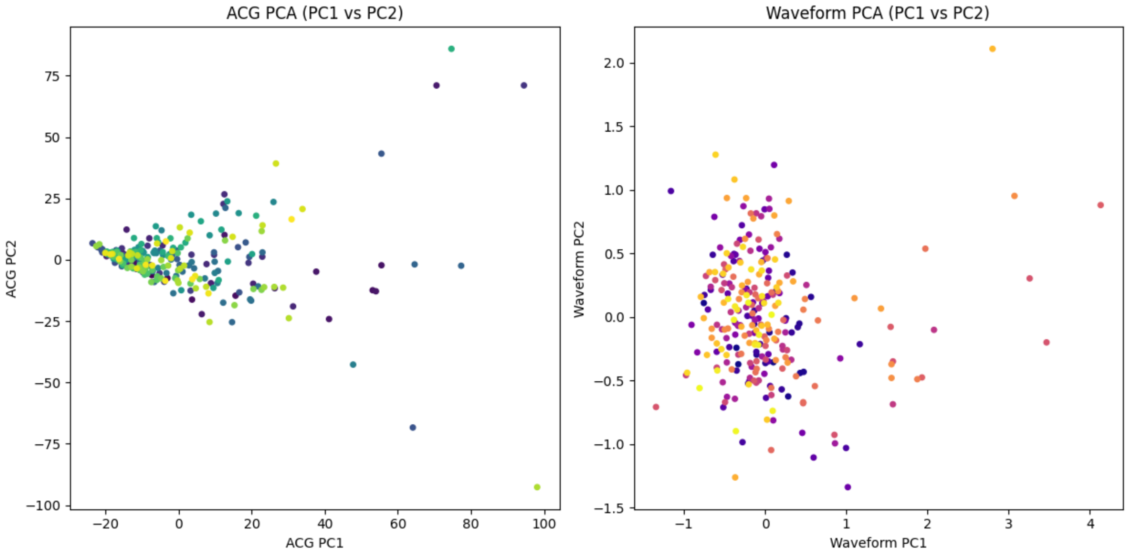


```
Number of valid clusters passing all filters: 268
Shape of normalized ACG subset: (268, 201)
Shape of collapsed waveforms: (268, 128)
Explained variance ratio (ACG PCA): [0.56357618 0.28218983]
Explained variance ratio (Waveform PCA): [0.436171   0.20883067]
Correlation between ACG PC1 and Waveform PC1: -0.1559
```

In [19]:

```
# Second Exploratory Iteration

import numpy as np
import matplotlib.pyplot as plt
from IPython.display import Markdown, display
import seaborn as sns

# For consistent plotting style
sns.set(context='notebook', style='whitegrid', font_scale=1.1)

#==================================================================
# (a) Display project status
```

```
#==================================================================
display(Markdown("""
**Exploratory Analysis - Iteration 2**

**Interpretation of Previous Results**
- From our initial PCA on ACGs and waveforms, we see that the first
two PCs capture most of the variance in each domain (56% + 28% for
ACG, and 44% + 21% for waveforms).
- There is a broad distribution of cells in PCA space, with no
immediate, obvious clustering.
- The correlation between ACG PC1 and waveform PC1 is quite low (-
0.156), suggesting that the primary axis of ACG variation does not
strongly co-vary with the primary axis of waveform variation.

**Current Working Hypotheses**
1. Multiple dimensions may be at play in the relationship between
ACG and waveform shape, and the main variance axes (PC1) might not
align in a simple way across ACG vs. waveform spaces.
2. Certain subgroups of neurons (e.g., bursting, narrow-spiking)
might occupy specific corners in the PCA space.
3. Brain region might explain some additional variance in both
waveform and ACG shape.

**Biological Interpretation**
- A sub-population of neurons could be fast-spiking inhibitory cells
(often narrower waveforms, more bursting ACG).
- Another sub-population might be regular-spiking excitatory cells
(broader waveforms, different ACG structure).

**Rationale for New Analysis**
To further explore how these PCs map to raw data, let's:
1. Pick a few representative cells spanning extremes in the joint
PCA space.
2. Visualize their raw ACG (in -200ms..+200ms) and their normalized
waveforms on the channel of max amplitude.
3. See whether the extremes of PCA indeed correspond to distinct
firing/waveform shapes.

"""))

#==================================================================
# (b) Perform the computational analysis
#==================================================================
# We assume we still have in memory (from previous step):
#   acg_norm          : shape [nValid, 201], normalized ACG from -
200..200 ms
#   collapsed_waveforms : shape [nValid, nTimes], normalized
waveforms
#   valid_indices
```

```
#   acg_scores, wave_scores : PCA-projected coordinates for ACG and
waveform
#   regions_valid     : brain region labels for the valid clusters
#
# For clarity, we define them as though they're in the namespace.

# We'll pick 5 representative clusters based on extremes in both ACG
PC1 and waveform PC1.
# Let's pick:
#   1) A cluster with max ACG PC1
#   2) A cluster with min ACG PC1
#   3) A cluster with max Wave PC1
#   4) A cluster with min Wave PC1
#   5) A cluster near the origin in both spaces

acg_pc1 = acg_scores[:,0]
wave_pc1 = wave_scores[:,0]

max_acg_idx = np.argmax(acg_pc1)
min_acg_idx = np.argmin(acg_pc1)
max_wav_idx = np.argmax(wave_pc1)
min_wav_idx = np.argmin(wave_pc1)

# find a cluster near (0,0) in (ACG PC1, Wave PC1) space
dists = np.sqrt(acg_pc1**2 + wave_pc1**2)
mid_idx = np.argmin(dists)

representative_indices = [max_acg_idx, min_acg_idx, max_wav_idx,
min_wav_idx, mid_idx]

# We'll retrieve the corresponding waveforms and ACG slices
rep_acgs = acg_norm[representative_indices]   # shape [5, 201]
rep_waves = collapsed_waveforms[representative_indices]  # shape [5,
nTimes]

# The time axis for waveforms (samples) - let's assume we have
nTimes=128, ~1 sample each.
wave_time = np.arange(rep_waves.shape[1])

# The lag axis for the ACG in -200..200 ms, in 2ms increments, so
length=201.
# If the midpoint is 0, then index 0 corresponds to -200 ms, etc.
acg_lags = np.linspace(-200, 200, 201)

#=================================================================
# (c) Generate a single matplotlib figure with subplots
#=================================================================
fig = plt.figure(figsize=(14, 8))
```

```
# Top row: scatter plots of ACG PCs and Wave PCs
ax1 = fig.add_subplot(2, 2, 1)
scatter1 = ax1.scatter(acg_scores[:,0], acg_scores[:,1],
                       c='gray', alpha=0.5, s=15, label='All
clusters')
ax1.set_title('ACG PCA (PC1 vs. PC2)')
ax1.set_xlabel('ACG PC1')
ax1.set_ylabel('ACG PC2')

# Circle our chosen points in the ACG PCA space
colors = ['red','blue','green','orange','purple']
for i, idx in enumerate(representative_indices):
    ax1.scatter(acg_scores[idx,0], acg_scores[idx,1],
                edgecolor='black', facecolor=colors[i], s=80,
label=f"Rep {i+1}" if i==0 else None)
ax1.legend(loc='best')

ax2 = fig.add_subplot(2, 2, 2)
scatter2 = ax2.scatter(wave_scores[:,0], wave_scores[:,1],
                       c='gray', alpha=0.5, s=15, label='All
clusters')
ax2.set_title('Waveform PCA (PC1 vs. PC2)')
ax2.set_xlabel('Wave PC1')
ax2.set_ylabel('Wave PC2')

# Circle the chosen points in wave PCA space
for i, idx in enumerate(representative_indices):
    ax2.scatter(wave_scores[idx,0], wave_scores[idx,1],
                edgecolor='black', facecolor=colors[i], s=80,
label=f"Rep {i+1}" if i==0 else None)
ax2.legend(loc='best')

# Bottom row: overlay each representative cluster's ACG and waveform
ax3 = fig.add_subplot(2, 2, 3)
for i in range(len(representative_indices)):
    ax3.plot(acg_lags, rep_acgs[i], color=colors[i], label=f"Rep
{i+1}")
ax3.set_title('Representative Clusters: ACG')
ax3.set_xlabel('Lag (ms)')
ax3.set_ylabel('Normalized count')
ax3.legend(loc='best')

ax4 = fig.add_subplot(2, 2, 4)
for i in range(len(representative_indices)):
    ax4.plot(wave_time, rep_waves[i], color=colors[i], label=f"Rep
{i+1}")
ax4.set_title('Representative Clusters: Waveform')
ax4.set_xlabel('Time samples')
ax4.set_ylabel('Norm. amplitude')
```

```
ax4.legend(loc='best')

plt.tight_layout()
plt.show()

#======================================================================
# (d) Print any numerical statistics that might help refine
hypothesis
#======================================================================
print("Representative clusters (indices in valid array):",
representative_indices)
print("ACG PC1 values for reps:", acg_pc1[representative_indices])
print("Wave PC1 values for reps:", wave_pc1[representative_indices])
print("Regions for reps:", regions_valid[representative_indices])

#======================================================================
# (e) Print the correlation coefficient from the previous step
#======================================================================
corr_coefficient = np.corrcoef(acg_pc1, wave_pc1)[0,1]
print(f"Correlation between ACG PC1 and Waveform PC1 (same as
before): {corr_coefficient:.4f}")

#END
```

**Exploratory Analysis - Iteration 2**

**Interpretation of Previous Results**

- From our initial PCA on ACGs and waveforms, we see that the first two PCs capture most of the variance in each domain (56% + 28% for ACG, and 44% + 21% for waveforms).
- There is a broad distribution of cells in PCA space, with no immediate, obvious clustering.
- The correlation between ACG PC1 and waveform PC1 is quite low (-0.156), suggesting that the primary axis of ACG variation does not strongly co-vary with the primary axis of waveform variation.

**Current Working Hypotheses**

1. Multiple dimensions may be at play in the relationship between ACG and waveform shape, and the main variance axes (PC1) might not align in a simple way across ACG vs. waveform spaces.
2. Certain subgroups of neurons (e.g., bursting, narrow-spiking) might occupy specific corners in the PCA space.
3. Brain region might explain some additional variance in both waveform and ACG shape.

**Biological Interpretation**

- A sub-population of neurons could be fast-spiking inhibitory cells (often narrower waveforms, more bursting ACG).
- Another sub-population might be regular-spiking excitatory cells (broader waveforms, different ACG structure).

**Rationale for New Analysis**
To further explore how these PCs map to raw data, let's:

1. Pick a few representative cells spanning extremes in the joint PCA space.
2. Visualize their raw ACG (in -200ms..+200ms) and their normalized waveforms on the channel of max amplitude.
3. See whether the extremes of PCA indeed correspond to distinct firing/waveform shapes.

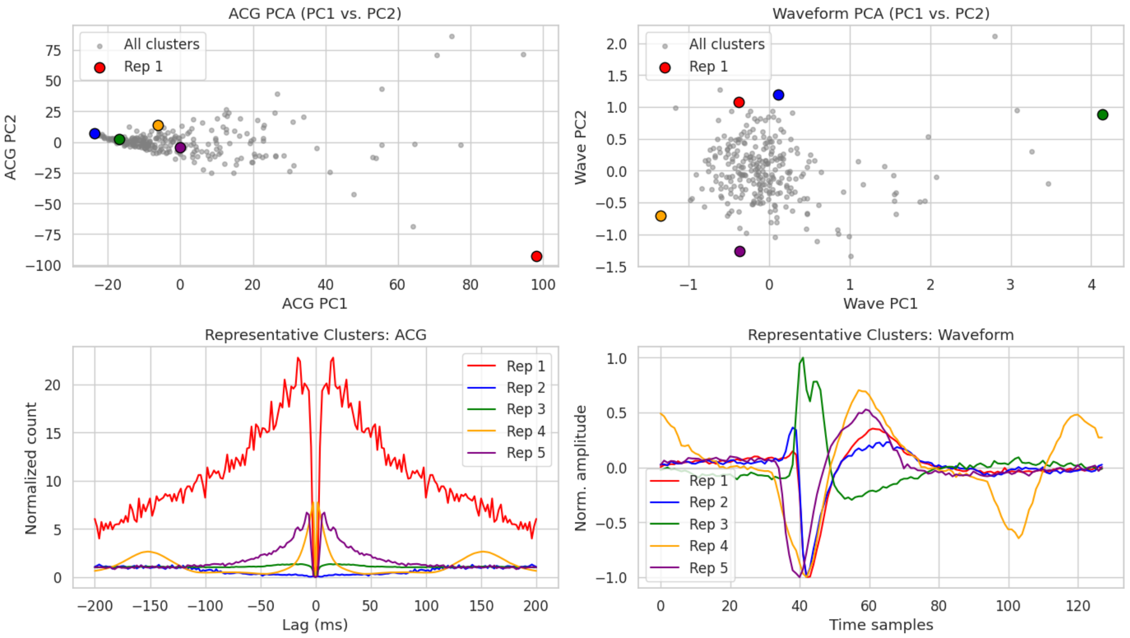


```
Representative clusters (indices in valid array): [234, 60, 154, 147, 219]
ACG PC1 values for reps: [ 98.14141162 -23.58492948 -16.93132133  -6.24891467  -0.13509235]
Wave PC1 values for reps: [-0.37866363  0.1097009   4.134194   -1.3457307  -0.3697854 ]
Regions for reps: ['CP' 'CP' 'SSp-m6a' 'SSp-m6a' 'EPd']
Correlation between ACG PC1 and Waveform PC1 (same as before): -0.1559
```

In [20]:

```
# Third Exploratory Iteration

import numpy as np
import matplotlib.pyplot as plt
from IPython.display import Markdown, display
import seaborn as sns
import pandas as pd
from collections import Counter

# For consistent plotting style
sns.set(context='notebook', style='whitegrid', font_scale=1.1)

#==================================================================
# (a) Display project status
```

```
#==========================================================================
display(Markdown("""
**Exploratory Analysis - Iteration 3**

**Interpretation of Previous Results**
- Our representative clusters demonstrate different types of
autocorrelogram (ACG) shapes (burstiness, broader or narrower peaks,
etc.) and waveform shapes (peak/trough widths).
- The correlation between ACG PC1 and Waveform PC1 remains small (-
0.156).
- We see large inter-individual diversity, suggesting that a simple
one-dimensional relationship may not explain ACG vs. waveform
variation.

**Current Working Hypotheses**
1. Some portion of the variance in ACG and waveform might be related
to brain region.
2. Different brain regions could show distinct distributions of ACG
or waveform PC values.

**Biological Interpretation**
- Brain regions often differ in their cell-type composition, which
could manifest in distinct firing patterns (ACGs) and/or waveform
shapes across regions.

**Rationale for New Analysis**
We will now color the scatter of ACG PC1 vs. Waveform PC1 by region,
then compare distributions across regions.
- This may reveal whether region helps explain any portion of the
variation in ACG or waveform PCs.
- We will also compute within-region correlation coefficients
between ACG PC1 and Waveform PC1, to see if the relationship varies
by region.
"""))

#==========================================================================
# (b) Perform the computational analysis
#==========================================================================
# We assume we have the following in memory:
#   valid_indices
#   acg_scores, wave_scores -> shape [nValid, 2]
#   regions_valid -> shape [nValid] (string array of region
acronyms)
# from the previous steps.

acg_pc1 = acg_scores[:, 0]
wave_pc1 = wave_scores[:, 0]

# Convert data to a pandas DataFrame for convenience
```

```
df = pd.DataFrame({
    'acg_pc1': acg_pc1,
    'wave_pc1': wave_pc1,
    'region': regions_valid
})

# Count how many cells per region
region_counts = Counter(df['region'])
# We might want to show only regions with at least a few cells.
Let's keep all for now.

# We'll compute within-region correlation (Pearson) between wave_pc1
and acg_pc1
unique_regions = np.unique(regions_valid)
region_corrs = {}
for reg in unique_regions:
    sub = df[df['region'] == reg]
    if len(sub) > 1:
        r = np.corrcoef(sub['acg_pc1'], sub['wave_pc1'])[0,1]
        region_corrs[reg] = r
    else:
        region_corrs[reg] = np.nan  # Only 1 cell => no correlation

#=====================================================================
# (c) Generate a single matplotlib figure with subplots
#=====================================================================
fig = plt.figure(figsize=(18, 6))

# Subplot 1: scatter of ACG PC1 vs Wave PC1, color by region
ax1 = fig.add_subplot(1, 3, 1)
# We'll pick a color palette
palette = sns.color_palette('tab20', n_colors=len(unique_regions))
region_to_color = {reg: palette[i % len(palette)] for i, reg in
enumerate(unique_regions)}

for reg in unique_regions:
    sub = df[df['region'] == reg]
    ax1.scatter(sub['acg_pc1'], sub['wave_pc1'],
                color=region_to_color[reg], label=reg, alpha=0.7,
s=25)

ax1.set_title('ACG PC1 vs Wave PC1 (Colored by Region)')
ax1.set_xlabel('ACG PC1')
ax1.set_ylabel('Wave PC1')
ax1.legend(loc='upper right', bbox_to_anchor=(1.35, 1.0), ncol=1,
frameon=True)

# Subplot 2: boxplot of ACG PC1 by region
ax2 = fig.add_subplot(1, 3, 2)
```

```
# For a nicer plot, let's show only regions with at least 5 cells
region_threshold = 5
keep_regions = [r for r in unique_regions if region_counts[r] >=
region_threshold]
df_box_acg = df[df['region'].isin(keep_regions)]

sns.boxplot(y='region', x='acg_pc1', data=df_box_acg, ax=ax2,
order=keep_regions, palette='tab20')
ax2.set_title('ACG PC1 by Region')
ax2.set_xlabel('ACG PC1')
ax2.set_ylabel('')

# Subplot 3: boxplot of Wave PC1 by region
ax3 = fig.add_subplot(1, 3, 3)
df_box_wave = df[df['region'].isin(keep_regions)]
sns.boxplot(y='region', x='wave_pc1', data=df_box_wave, ax=ax3,
order=keep_regions, palette='tab20')
ax3.set_title('Wave PC1 by Region')
ax3.set_xlabel('Wave PC1')
ax3.set_ylabel('')

plt.tight_layout()
plt.show()

#=====================================================================
# (d) Print numeric statistics
#=====================================================================
print("Number of unique regions in valid set:", len(unique_regions))
print("Region counts:", region_counts)
print("Within-region correlation between ACG PC1 and Wave PC1:")
for reg in unique_regions:
    print(f"  {reg}: {region_corrs[reg]:.3f}" if not
np.isnan(region_corrs[reg])
          else f"  {reg}: (not enough cells)")

#=====================================================================
# (e) Print the overall correlation coefficient again
#=====================================================================
overall_corr = np.corrcoef(df['acg_pc1'], df['wave_pc1'])[0,1]
print(f"\nOverall correlation (ACG PC1 vs Wave PC1) across all
regions: {overall_corr:.4f}")

#END
```

**Exploratory Analysis - Iteration 3**

**Interpretation of Previous Results**

- Our representative clusters demonstrate different types of autocorrelogram (ACG) shapes (burstiness, broader or narrower peaks, etc.) and waveform shapes (peak/trough widths).

- The correlation between ACG PC1 and Waveform PC1 remains small (-0.156).
- We see large inter-individual diversity, suggesting that a simple one-dimensional relationship may not explain ACG vs. waveform variation.

**Current Working Hypotheses**

1. Some portion of the variance in ACG and waveform might be related to brain region.
2. Different brain regions could show distinct distributions of ACG or waveform PC values.

**Biological Interpretation**

- Brain regions often differ in their cell-type composition, which could manifest in distinct firing patterns (ACGs) and/or waveform shapes across regions.

**Rationale for New Analysis**
We will now color the scatter of ACG PC1 vs. Waveform PC1 by region, then compare distributions across regions.

- This may reveal whether region helps explain any portion of the variation in ACG or waveform PCs.
- We will also compute within-region correlation coefficients between ACG PC1 and Waveform PC1, to see if the relationship varies by region.

```
<ipython-input-20-9e1c243d4f44>:99: FutureWarning:

Passing `palette` without assigning `hue` is deprecated and will be removed in v0.14.0. Assign the `y` variable to `hue` and set `legend=False` for the same effect.

  sns.boxplot(y='region', x='acg_pc1', data=df_box_acg, ax=ax2, order=keep_regions, palette='tab20')
<ipython-input-20-9e1c243d4f44>:107: FutureWarning:

Passing `palette` without assigning `hue` is deprecated and will be removed in v0.14.0. Assign the `y` variable to `hue` and set `legend=False` for the same effect.

  sns.boxplot(y='region', x='wave_pc1', data=df_box_wave, ax=ax3, order=keep_regions, palette='tab20')
```

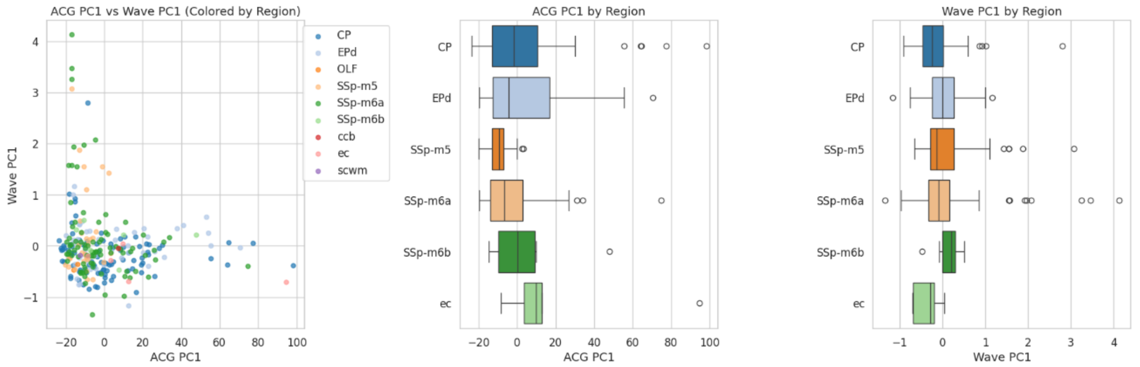


```
Number of unique regions in valid set: 9
```

```
Region counts: Counter({'CP': 94, 'SSp-m6a': 82, 'EPd': 48, 'SSp-m5': 29, 'SSp-m6b': 6, 'ec': 5, 'OLF': 2, 'ccb': 1, 'scwm': 1})
Within-region correlation between ACG PC1 and Wave PC1:
  CP: -0.103
  EPd: 0.074
  OLF: 1.000
  SSp-m5: 0.057
  SSp-m6a: -0.278
  SSp-m6b: -0.111
  ccb: (not enough cells)
  ec: -0.626
  scwm: (not enough cells)

Overall correlation (ACG PC1 vs Wave PC1) across all regions: -0.1559
```

In [21]:

```
# Fourth Exploratory Iteration

import numpy as np
import matplotlib.pyplot as plt
import seaborn as sns
from IPython.display import Markdown, display
from umap import UMAP  # make sure the user has "pip install umap-learn"
import pandas as pd

sns.set(context='notebook', style='whitegrid', font_scale=1.1)

#=====================================================================
# (a) Display project status
#=====================================================================
display(Markdown("""
**Exploratory Analysis – Iteration 4**

**Interpretation of Previous Results**
- We found that different brain regions show distinct distributions in both ACG PC1 and Wave PC1.
- The overall correlation between ACG PC1 and Wave PC1 remains modest (-0.156).
- Within-region correlations vary but remain low, suggesting no strong linear relationship in any single region.

**Current Working Hypotheses**
1. A higher-dimensional relationship between ACG shape and waveform shape may exist that is not well-captured by a single PC.
2. Non-linear embeddings (e.g., UMAP) might better reveal clusters or manifold structure shared between ACG and waveform features.
3. Brain-region differences might become more apparent (or less confounded) in a joint embedding.
```

```
**Biological Interpretation**
- Certain cell types (fast-spiking, regular spiking, etc.) likely
occur in multiple regions, but in different proportions,
complicating the interpretation of region-based trends.
- Non-linear embeddings could group neurons by common morphological
or firing properties that cut across region boundaries.

**Rationale for New Analysis**
We will:
1. Combine the normalized ACG features (from -200..200 ms) and the
normalized waveform features (collapsed onto the max channel) into
one feature vector per cell.
2. Use UMAP to embed these combined features into a 2D space.
3. Color by region to see if region-based clusters emerge, and look
for potential structure relating ACG and waveform shapes.
"""))

#==================================================================
# (b) Perform the computational analysis
#==================================================================
# Assume we still have in memory:
#   acg_norm : shape [nValid, 201], normalized ACG from -200..200ms
#   collapsed_waveforms : shape [nValid, nTimes], normalized
waveforms
#   regions_valid : shape [nValid], region acronyms
# from prior steps.

# We'll combine ACG + waveform features for each cell into one
vector
combined_features = np.concatenate([acg_norm, collapsed_waveforms],
axis=1)
# shape is [nValid, 201 + nTimes] = [nValid, 201 + 128] = [nValid,
329]

# Instantiate and run UMAP
umap_model = UMAP(n_neighbors=15, min_dist=0.1, n_components=2,
random_state=42)
umap_coords = umap_model.fit_transform(combined_features)  # shape
[nValid, 2]

#==================================================================
# (c) Generate a single matplotlib figure with subplots
#==================================================================
fig = plt.figure(figsize=(10, 8))
ax = fig.add_subplot(1, 1, 1)

regions_unique = np.unique(regions_valid)
palette = sns.color_palette('tab20', n_colors=len(regions_unique))
```

```
region_color_map = {reg: palette[i % len(palette)] for i, reg in
enumerate(regions_unique)}

for reg in regions_unique:
    idx = np.where(regions_valid == reg)[0]
    ax.scatter(umap_coords[idx, 0], umap_coords[idx, 1],
               c=[region_color_map[reg]]*len(idx),
               alpha=0.7, label=reg, s=25)

ax.set_title("UMAP of Combined ACG + Waveform Features")
ax.set_xlabel("UMAP Dimension 1")
ax.set_ylabel("UMAP Dimension 2")
ax.legend(loc='upper right', bbox_to_anchor=(1.35, 1.0), ncol=1,
frameon=True)
plt.tight_layout()
plt.show()

#==================================================================
# (d) Print numeric statistics that might help refine hypotheses
#==================================================================
# We'll look at the variance in the new 2D embedding, though UMAP
doesn't provide
# a direct "explained variance" metric. Instead, let's consider how
well it separates
# regions. One measure could be silhouette score by region, though
that is a bit of
# a "cluster metric." We'll compute it as a rough guide.

from sklearn.metrics import silhouette_score

# We'll encode each region as a label
region_labels = pd.Categorical(regions_valid).codes  # integer
labels
if len(np.unique(region_labels)) > 1:
    sil_score = silhouette_score(umap_coords, region_labels)
else:
    sil_score = np.nan

print(f"UMAP feature dimension: {combined_features.shape}")
print(f"Computed 2D UMAP coords shape: {umap_coords.shape}")
print(f"Approximate silhouette score by region: {sil_score:.3f}")

#==================================================================
# (e) We won't compute a single correlation coefficient here,
#     since UMAP is a non-linear embedding. We'll simply print a
note.
#==================================================================
print("Note: UMAP is non-linear, so there's no simple correlation
metric analogous to PCA PC1 vs PC1.")
```

```
print("We'll use the silhouette score as a rough gauge of region
separability in 2D space.")
#END
```

**Exploratory Analysis – Iteration 4**

**Interpretation of Previous Results**

- We found that different brain regions show distinct distributions in both ACG PC1 and Wave PC1.
- The overall correlation between ACG PC1 and Wave PC1 remains modest (-0.156).
- Within-region correlations vary but remain low, suggesting no strong linear relationship in any single region.

**Current Working Hypotheses**

1. A higher-dimensional relationship between ACG shape and waveform shape may exist that is not well-captured by a single PC.
2. Non-linear embeddings (e.g., UMAP) might better reveal clusters or manifold structure shared between ACG and waveform features.
3. Brain-region differences might become more apparent (or less confounded) in a joint embedding.

**Biological Interpretation**

- Certain cell types (fast-spiking, regular spiking, etc.) likely occur in multiple regions, but in different proportions, complicating the interpretation of region-based trends.
- Non-linear embeddings could group neurons by common morphological or firing properties that cut across region boundaries.

**Rationale for New Analysis**
We will:

1. Combine the normalized ACG features (from -200..200 ms) and the normalized waveform features (collapsed onto the max channel) into one feature vector per cell.
2. Use UMAP to embed these combined features into a 2D space.
3. Color by region to see if region-based clusters emerge, and look for potential structure relating ACG and waveform shapes.

```
/usr/local/lib/python3.11/dist-
packages/sklearn/utils/deprecation.py:151: FutureWarning:
'force_all_finite' was renamed to 'ensure_all_finite' in 1.6 and
will be removed in 1.8.
  warnings.warn(
/usr/local/lib/python3.11/dist-packages/umap/umap_.py:1952:
UserWarning: n_jobs value 1 overridden to 1 by setting random_state.
Use no seed for parallelism.
  warn(
```

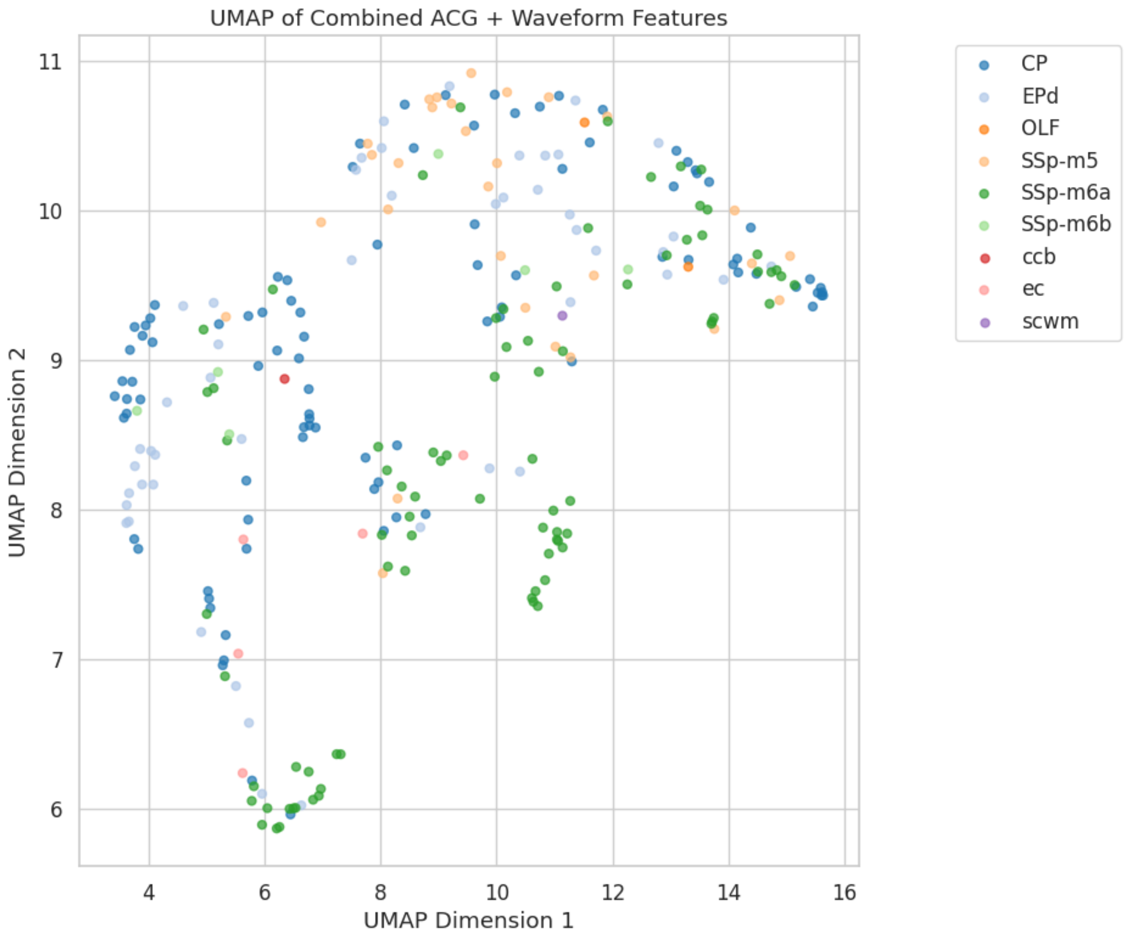


```
UMAP feature dimension: (268, 329)
Computed 2D UMAP coords shape: (268, 2)
Approximate silhouette score by region: -0.564
Note: UMAP is non-linear, so there's no simple correlation metric
analogous to PCA PC1 vs PC1.
We'll use the silhouette score as a rough gauge of region
separability in 2D space.
```

In [22]:

```
# Fifth Exploratory Iteration

import numpy as np
import matplotlib.pyplot as plt
import seaborn as sns
from IPython.display import Markdown, display
from sklearn.decomposition import PCA
from sklearn.cross_decomposition import CCA

sns.set(context='notebook', style='whitegrid', font_scale=1.1)

#==================================================================
# (a) Display project status
#==================================================================
display(Markdown("""
```

```
**Exploratory Analysis – Iteration 5**

**Interpretation of Previous Results**
- The UMAP embedding on combined ACG + waveform features does not
show strong region-based clustering (silhouette score: ~-0.56).
- This suggests that, in this dataset, cell firing/waveform
properties do not strictly segregate by region, or at least not in a
way UMAP can easily separate.

**Current Working Hypotheses**
1. Even though region-based clustering is weak, there may be a
linear relationship between ACG and waveform features beyond the
first principal components.
2. Canonical Correlation Analysis (CCA) could reveal linear
combinations of ACG features and waveform features that are
maximally correlated, potentially higher than the -0.16 observed
between PC1s.

**Biological Interpretation**
- A single principal component might not capture subtle aspects of
spike timing or waveform shape that do correlate across modalities.
- CCA can find latent, linearly-related dimensions that might
reflect, for instance, “burstiness plus spike shape,” or “regular
spiking plus narrower waveforms.”

**Rationale for New Analysis**
Here we will:
1. Reduce dimensionality of both the ACG and waveform feature sets
(to mitigate rank issues).
2. Use CCA on these reduced dimensions to find canonical variates
that are maximally correlated.
3. Plot the first canonical variate from the ACG side vs. the first
canonical variate from the waveform side, color by region, and
report the correlation.
"""))

#=================================================================
# (b) Perform the computational analysis
#=================================================================
# We'll assume we have the following from prior steps:
#   acg_norm : shape [nValid, 201]   (normalized ACG from -200..200
ms)
#   collapsed_waveforms : shape [nValid, nTimes]   (normalized
waveforms)
#   regions_valid : shape [nValid]  (string array of region
acronyms)
#   valid_indices : indices of all valid clusters
```

```
# 1) We first reduce each set (ACG, wave) to 10 PCs to avoid
dimensionality issues in CCA.
n_components_pca = 10

# ACG PCA
acg_pca = PCA(n_components=n_components_pca)
acg_pca_scores = acg_pca.fit_transform(acg_norm)  # shape [nValid,
10]

# Waveform PCA
wave_pca = PCA(n_components=n_components_pca)
wave_pca_scores = wave_pca.fit_transform(collapsed_waveforms)  #
shape [nValid, 10]

# 2) Perform CCA on these PCA-reduced features
cca = CCA(n_components=2)  # let's get the first two canonical
variates
cca.fit(acg_pca_scores, wave_pca_scores)

acg_cv, wave_cv = cca.transform(acg_pca_scores, wave_pca_scores)
# shape of each is [nValid, 2], representing canonical variates

# We'll look at the correlation of the first canonical dimension
cca_corr = np.corrcoef(acg_cv[:,0], wave_cv[:,0])[0,1]

#=================================================================
# (c) Generate a single matplotlib figure with subplots
#=================================================================
fig = plt.figure(figsize=(9, 6))
ax = fig.add_subplot(1, 1, 1)

# We'll color points by region
regions_unique = np.unique(regions_valid)
palette = sns.color_palette('tab20', n_colors=len(regions_unique))
region_color_map = {reg: palette[i % len(palette)] for i, reg in
enumerate(regions_unique)}

for reg in regions_unique:
    idx = np.where(regions_valid == reg)[0]
    ax.scatter(acg_cv[idx, 0], wave_cv[idx, 0],
               c=[region_color_map[reg]]*len(idx),
               alpha=0.7, s=25, label=reg)

ax.set_title('First Canonical Variates (ACG vs. Waveform)')
ax.set_xlabel('ACG Canonical Variate 1')
ax.set_ylabel('Wave Canonical Variate 1')
ax.legend(loc='upper right', bbox_to_anchor=(1.4, 1.0),
frameon=True)
plt.tight_layout()
```

```
plt.show()

#==================================================================
# (d) Print numeric statistics
#==================================================================
print(f"Shape of ACG PCA scores: {acg_pca_scores.shape}")
print(f"Shape of Waveform PCA scores: {wave_pca_scores.shape}")
print("Explained variance ratio (ACG PCA, first 10 PCs):",
acg_pca.explained_variance_ratio_)
print("Explained variance ratio (Waveform PCA, first 10 PCs):",
wave_pca.explained_variance_ratio_)

#==================================================================
# (e) Print the correlation of the first canonical variate
#==================================================================
print(f"Correlation of Canonical Variate 1 (ACG vs Waveform):
{cca_corr:.4f}")

#END
```

**Exploratory Analysis – Iteration 5**

**Interpretation of Previous Results**

- The UMAP embedding on combined ACG + waveform features does not show strong region-based clustering (silhouette score: ~-0.56).
- This suggests that, in this dataset, cell firing/waveform properties do not strictly segregate by region, or at least not in a way UMAP can easily separate.

**Current Working Hypotheses**

1. Even though region-based clustering is weak, there may be a linear relationship between ACG and waveform features beyond the first principal components.
2. Canonical Correlation Analysis (CCA) could reveal linear combinations of ACG features and waveform features that are maximally correlated, potentially higher than the -0.16 observed between PC1s.

**Biological Interpretation**

- A single principal component might not capture subtle aspects of spike timing or waveform shape that do correlate across modalities.
- CCA can find latent, linearly-related dimensions that might reflect, for instance, "burstiness plus spike shape," or "regular spiking plus narrower waveforms."

**Rationale for New Analysis**
Here we will:

1. Reduce dimensionality of both the ACG and waveform feature sets (to mitigate rank issues).
2. Use CCA on these reduced dimensions to find canonical variates that are maximally correlated.
3. Plot the first canonical variate from the ACG side vs. the first canonical variate from the waveform side, color by region, and report the correlation.

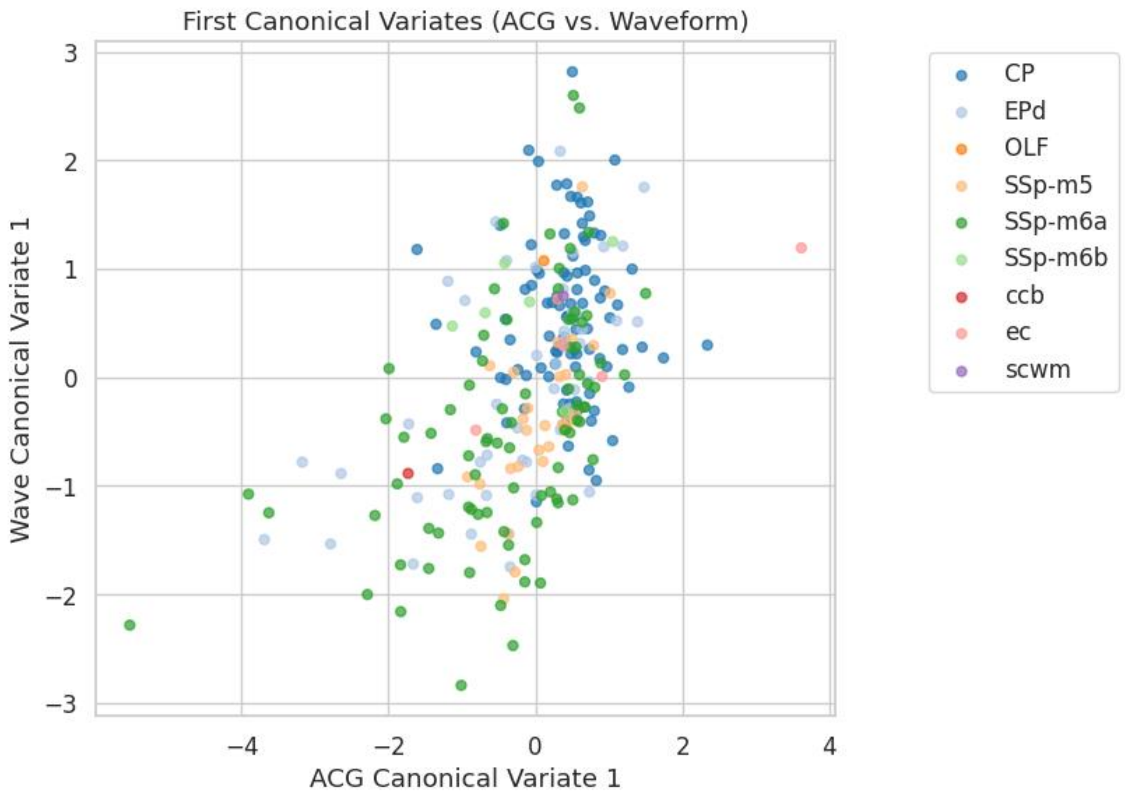


```
Shape of ACG PCA scores: (268, 10)
Shape of Waveform PCA scores: (268, 10)
Explained variance ratio (ACG PCA, first 10 PCs): [0.56357618
0.28218983 0.07375176 0.04525022 0.01082617 0.00684133
 0.00309944 0.00241195 0.00145    0.00106539]
Explained variance ratio (Waveform PCA, first 10 PCs): [0.436171
0.20883067 0.13070191 0.09373006 0.0347827  0.0281242
 0.01688985 0.01098075 0.0091475  0.00503077]
Correlation of Canonical Variate 1 (ACG vs Waveform): 0.4927
```

In [23]:

```
# Confirmatory Phase - Iteration 1

import numpy as np
import matplotlib.pyplot as plt
from IPython.display import Markdown, display
import seaborn as sns

# Make sure we've got consistent styling
sns.set(context='notebook', style='whitegrid', font_scale=1.1)

#==================================================================
# (a) Display project status
#==================================================================
display(Markdown("""
## Confirmatory Phase – Iteration 1
```

```
**Interpretation of Exploratory Results**
- The exploratory analyses suggest that firing-pattern features in
the ACG and shape features in the spike waveform can correlate along
certain dimensions (CCA gave us ~0.49 correlation on the first
canonical variate).
- We now transition to a more direct, confirmatory approach: we will
define **single scalar statistics** from the ACG and from the
waveform for each neuron, and test their correlation.

**Current Working Hypotheses**
1. Neurons that are more "burst-prone" (i.e., higher short-lag ACG
ratio) may also exhibit narrower waveforms.
2. By focusing on a short-lag ratio in the ACG and a trough-peak
timing in the waveform, we may see a moderate correlation.

**Biological Interpretation**
- Burstiness often corresponds to certain interneuron types, which
might have narrower waveforms.
- Regular-spiking excitatory cells might be less bursty and have
broader waveforms.

**Plan**
- Define a short-lag ratio from the ACG: \\
  \\( \mathrm{ACG\_stat} = \frac{\sum_{-10 \mathrm{ms}}^{+10
\mathrm{ms}} \mathrm{ACG}(t)}{\sum_{100 \mathrm{ms}}^{200
\mathrm{ms}} \mathrm{ACG}(t)}} \\).
- Define a waveform trough-peak interval: the difference (in
samples) between the most negative point and the most positive point
within each neuron's normalized waveform (on the max amplitude
channel).
- We then compute the correlation between these two statistics,
color points by region, and circle a handful of example neurons for
closer inspection of their raw ACG and waveform.

"""))

#==================================================================
# (b) Perform the computational analysis
#==================================================================

# We assume the following arrays in memory (from prior steps):
#   acg_norm: shape [nValid, 201] – normalized ACG from -200..+200ms
in 2ms bins
#   collapsed_waveforms: shape [nValid, nTimes=128] – normalized
waveforms on largest channel
#   regions_valid: shape [nValid] – region strings
#   valid_indices: used to identify these clusters among the full
dataset
#   acgs: shape [nClusters, 501], the original ACG if needed
```

```
#   waveforms.templates: shape [nClusters, nChannels, nTimes] for
original waveforms
#   clusters.acronym: shape [nClusters], for region
#
# For the short-lag ratio, we need index ranges:
#   -10..+10 ms => each bin is 2 ms, and our subset acg_norm is from
-200..+200 ms => total 201 bins
#   The center bin is index = 100 (0 ms).
#   So the short-lag region is from index 100 - 5 bins to 100 + 5
bins (11 total bins).
short_lag_inds = np.arange(95, 106)  # from -10ms to +10ms in steps
of 2ms
#   The far-lag region is from +100ms..+200ms => that is from index
150..200 in the [0..200] indexing.
#   Actually from +100 ms => index=100 + 50=150 to +200 ms =>
index=200 => so range(150, 201)
far_lag_inds = np.arange(150, 201)

acg_stat = np.sum(acg_norm[:, short_lag_inds], axis=1) / (1e-12 +
np.sum(acg_norm[:, far_lag_inds], axis=1))

# For the waveform statistic, let's define "trough_peak_interval":
#   1) Identify the trough index (most negative value)
#   2) Identify the peak index (most positive value)
#   3) difference in sample indices (peakIdx - troughIdx)
# We do this for each row of collapsed_waveforms.

wave_stat = []
for i in range(collapsed_waveforms.shape[0]):
    wv = collapsed_waveforms[i]
    trough_idx = np.argmin(wv)
    peak_idx = np.argmax(wv)
    wave_stat.append(peak_idx - trough_idx)
wave_stat = np.array(wave_stat)

# We'll compute the Pearson correlation
corr_coef = np.corrcoef(acg_stat, wave_stat)[0,1]

#==================================================================
# (c) Generate a single figure
#==================================================================
fig = plt.figure(figsize=(14, 8))

# Top-left: scatter of the two stats, color by region
ax1 = fig.add_subplot(2, 2, 1)
unique_regions = np.unique(regions_valid)
palette = sns.color_palette('tab10', n_colors=len(unique_regions))
region_to_color = {r: palette[i % len(palette)] for i, r in
enumerate(unique_regions)}
```

```
for r in unique_regions:
    idx = np.where(regions_valid == r)[0]
    ax1.scatter(acg_stat[idx], wave_stat[idx],
color=region_to_color[r], alpha=0.7, label=r, s=20)

ax1.set_xlabel('Short-lag ratio (ACG)')
ax1.set_ylabel('Trough-to-peak interval (samples)')
ax1.set_title('ACG vs Waveform Statistic')
ax1.legend(loc='best', prop={'size': 7})

# We'll circle ~6 example neurons: pick extremes + random
#   1) highest ACG stat
#   2) lowest ACG stat
#   3) highest wave stat
#   4) lowest wave stat
#   2 random
inds_sorted_acg = np.argsort(acg_stat)
inds_sorted_wav = np.argsort(wave_stat)

ex_inds = [
    inds_sorted_acg[-1],
    inds_sorted_acg[0],
    inds_sorted_wav[-1],
    inds_sorted_wav[0]
]
# Add 2 random from the middle
nMid = len(acg_stat) // 2
mid_slice = np.arange(nMid-10, nMid+10)
ex_inds += list(np.random.choice(mid_slice, size=2, replace=False))

ex_inds = list(dict.fromkeys(ex_inds))  # remove duplicates
colors_examples = ['red','blue','orange','green','magenta','purple']

for i, exi in enumerate(ex_inds):
    ax1.scatter(acg_stat[exi], wave_stat[exi],
                facecolors='none', edgecolors=colors_examples[i],
s=100, linewidths=2)

# Next, let's plot their ACG and waveforms
# We'll need to retrieve the raw subsets from acg_norm and
collapsed_waveforms
acg_lags = np.linspace(-200, 200, 201)
wave_time = np.arange(collapsed_waveforms.shape[1])

ax2 = fig.add_subplot(2, 2, 2)
for i, exi in enumerate(ex_inds):
    ax2.plot(acg_lags, acg_norm[exi], color=colors_examples[i],
label=f"Cell {exi}")
```

```
ax2.set_title('Example Cells: ACG')
ax2.set_xlabel('Lag (ms)')
ax2.set_ylabel('Normalized count')
ax2.legend(loc='best', prop={'size': 6})

ax3 = fig.add_subplot(2, 1, 2)
for i, exi in enumerate(ex_inds):
    ax3.plot(wave_time, collapsed_waveforms[exi],
color=colors_examples[i], label=f"Cell {exi}")
ax3.set_title('Example Cells: Waveforms')
ax3.set_xlabel('Time (samples)')
ax3.set_ylabel('Norm. amplitude')
ax3.legend(loc='best', prop={'size': 6})

plt.tight_layout()
plt.show()

#==================================================================
# (d) Print any statistics for diagnosing the fit
#==================================================================
print(f"Number of cells in analysis: {len(acg_stat)}")
print("ACG short-lag ratio: mean={:.3f},
std={:.3f}".format(np.mean(acg_stat), np.std(acg_stat)))
print("Wave trough-peak interval: mean={:.2f} samples,
std={:.2f}".format(np.mean(wave_stat), np.std(wave_stat)))

#==================================================================
# (e) Print correlation coefficient
#==================================================================
print(f"Correlation between short-lag ratio and trough-peak
interval: {corr_coef:.4f}")

#END
```

### Confirmatory Phase – Iteration 1

**Interpretation of Exploratory Results**

- The exploratory analyses suggest that firing-pattern features in the ACG and shape features in the spike waveform can correlate along certain dimensions (CCA gave us ~0.49 correlation on the first canonical variate).
- We now transition to a more direct, confirmatory approach: we will define **single scalar statistics** from the ACG and from the waveform for each neuron, and test their correlation.

**Current Working Hypotheses**

1. Neurons that are more "burst-prone" (i.e., higher short-lag ACG ratio) may also exhibit narrower waveforms.
2. By focusing on a short-lag ratio in the ACG and a trough-peak timing in the waveform, we may see a moderate correlation.

**Biological Interpretation**

- Burstiness often corresponds to certain interneuron types, which might have narrower waveforms.
- Regular-spiking excitatory cells might be less bursty and have broader waveforms.

**Plan**

- Define a short-lag ratio from the ACG: \ ( \mathrm{ACG_stat} =

rac{\sum{-10 \mathrm{ms}}^{+10 \mathrm{ms}} \mathrm{ACG}(t)}{\sum{100 \mathrm{ms}}^{200 \mathrm{ms}} \mathrm{ACG}(t)}} ).

- Define a waveform trough-peak interval: the difference (in samples) between the most negative point and the most positive point within each neuron's normalized waveform (on the max amplitude channel).
- We then compute the correlation between these two statistics, color points by region, and circle a handful of example neurons for closer inspection of their raw ACG and waveform.

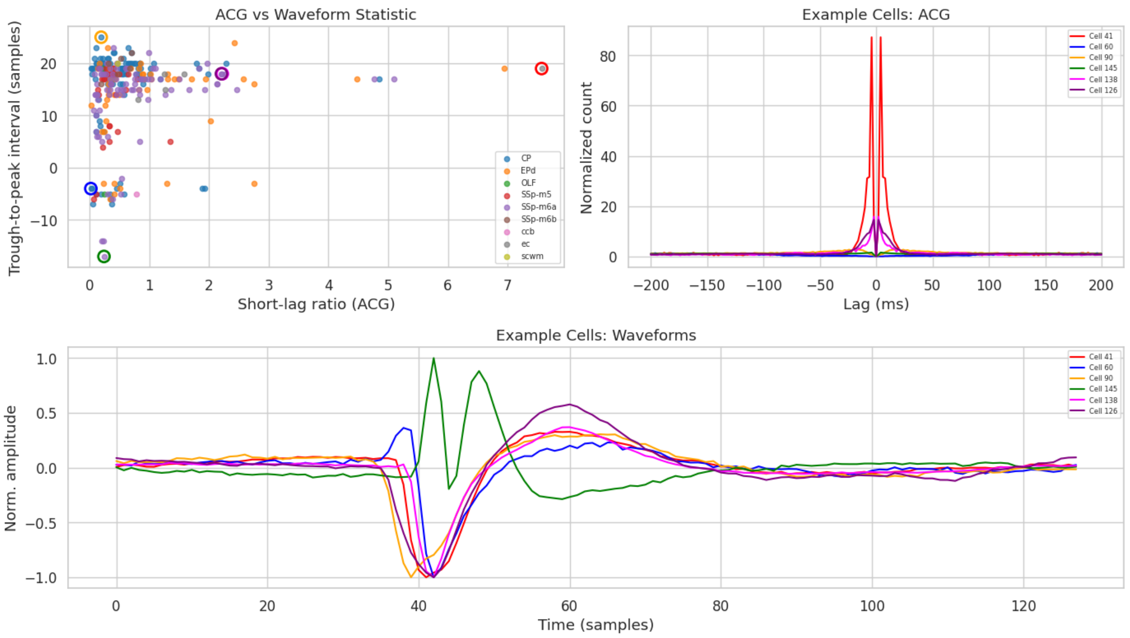


```
Number of cells in analysis: 268
ACG short-lag ratio: mean=0.740, std=0.947
Wave trough-peak interval: mean=14.12 samples, std=8.09
Correlation between short-lag ratio and trough-peak interval: 0.1277
```

In [24]:

```
# Confirmatory Phase - Iteration 2

import numpy as np
import matplotlib.pyplot as plt
import seaborn as sns
from IPython.display import Markdown, display

sns.set(context='notebook', style='whitegrid', font_scale=1.1)

#==================================================================
# (a) Display project status
#==================================================================
display(Markdown(r"""
## Confirmatory Phase – Iteration 2

**Interpretation of Previous Results**
- We used a short-lag ACG ratio and the trough-to-peak sample
difference in the waveform as our two metrics.
```

- The correlation was about 0.13, indicating a fairly weak linear relationship.

**Potential Reasons**
1. The short-lag ratio might be a bit coarse and could saturate for strongly bursting neurons.
2. The trough-to-peak *interval* alone might not fully capture broader vs. narrower waveforms.

**Revised Hypothesis**
- Perhaps a measure of ACG *peak width* at half-height (which might reflect burst width) will correlate more strongly with a measure of waveform *width* at half-trough (often used to distinguish fast- vs. regular-spiking cells).

**Plan**
- We will define two new statistics:
  1. **ACG Half-Width**: In the normalized ACG around 0 ms, find the peak near t=0, then measure how many ms it takes to go from the peak down to half its height on both sides, and sum that range.
  2. **Waveform Half-Width**: For the collapsed, normalized waveform, find the trough's minimum amplitude, then measure how many samples it takes on either side of that trough to rise to *half* of that amplitude. (Equivalently, "full-width at half-trough.")

We'll then plot these two new metrics, color by region, and quantify their correlation.
"""))

```
#==================================================================
# (b) Perform the computational analysis
#==================================================================
# Assumes we still have in memory:
#   acg_norm : shape [nValid, 201], normalized from -200..+200 ms in 2ms bins
#   collapsed_waveforms : shape [nValid, 128], normalized waveforms
#   valid_indices, regions_valid
#
# We'll define the new metrics:

acg_lags = np.linspace(-200, 200, 201)  # in ms
dt_acg = 2.0  # each bin is 2 ms

def compute_acg_halfwidth(acg_row, lag_ms, center_tolerance=15):
    """
    1) Identify the main peak around t=0 by looking in ±center_tolerance ms.
    2) Find the maximum in that range.
    3) Find the time range where the ACG is above half that maximum.
```

```
       The difference in ms is the half-width.
    If the peak is extremely sharp or half-max not found, returns 0
or minimal width.
    """
    # 1) Restrict to ±center_tolerance ms around 0
    mask_center = (lag_ms >= -center_tolerance) & (lag_ms <=
center_tolerance)
    sub_acg = acg_row[mask_center]
    sub_lags = lag_ms[mask_center]
    if len(sub_acg) < 1:
        return 0.0

    # 2) Max in that range
    peak_val = np.max(sub_acg)
    if peak_val < 1e-12:
        return 0.0
    half_val = 0.5 * peak_val

    # 3) Indices in sub_acg that exceed half_val
    above_half_mask = sub_acg >= half_val
    if not np.any(above_half_mask):
        return 0.0

    # get min & max index in sub_lags that are above half
    i_min = np.where(above_half_mask)[0][0]
    i_max = np.where(above_half_mask)[0][-1]

    # the difference in sub_lags at i_max - i_min is the halfwidth
    width = sub_lags[i_max] - sub_lags[i_min]
    return max(width, 0.0)

acg_halfwidth = np.array([compute_acg_halfwidth(acg_row, acg_lags)
                          for acg_row in acg_norm])


def compute_waveform_halfwidth(wave):
    """
    1) Identify the trough (most negative sample).
    2) The trough value = wave[trough_idx].
       Let half_val = trough_value / 2 (a negative number with
smaller magnitude).
    3) Find the points around the trough where the waveform crosses
half_val
       (on the left and right side). The difference in samples is
the half-width.
    4) Return as a positive number (in samples).
    If trough_value >= 0 (some strange wave), return 0.
    """
    trough_idx = np.argmin(wave)
```

```
    trough_val = wave[trough_idx]
    if trough_val >= 0:
        return 0.0

    half_val = trough_val / 2.0  # note trough_val is negative =>
half_val is also negative, but "less negative"

    # find left crossing
    left_idx = trough_idx
    while left_idx > 0 and wave[left_idx] < half_val:
        left_idx -= 1
    # find right crossing
    right_idx = trough_idx
    while right_idx < len(wave)-1 and wave[right_idx] < half_val:
        right_idx += 1

    # difference in sample indices
    halfwidth_samples = right_idx - left_idx
    if halfwidth_samples < 0:
        halfwidth_samples = 0
    return float(halfwidth_samples)

wave_halfwidth = np.array([compute_waveform_halfwidth(wv) for wv in
collapsed_waveforms])

# Now compute correlation
corr_coef = np.corrcoef(acg_halfwidth, wave_halfwidth)[0,1]

#==================================================================
# (c) Generate a single figure
#==================================================================
fig = plt.figure(figsize=(14, 8))

ax1 = fig.add_subplot(2, 2, 1)
unique_regions = np.unique(regions_valid)
palette = sns.color_palette('tab10', n_colors=len(unique_regions))
region_to_color = {r: palette[i % len(palette)] for i, r in
enumerate(unique_regions)}

for r in unique_regions:
    idx = np.where(regions_valid == r)[0]
    ax1.scatter(acg_halfwidth[idx], wave_halfwidth[idx],
                color=region_to_color[r], alpha=0.7, label=r, s=20)

ax1.set_xlabel('ACG Half-Width (ms)')
ax1.set_ylabel('Waveform Half-Width (samples)')
ax1.set_title('New ACG vs Waveform Statistic')
ax1.legend(loc='best', prop={'size': 7})
```

```
# Circle a few examples again
ex_indices = []
ex_indices.append(np.argmax(acg_halfwidth))  # largest ACG halfwidth
ex_indices.append(np.argmin(acg_halfwidth))  # smallest
ex_indices.append(np.argmax(wave_halfwidth)) # largest wave
halfwidth
ex_indices.append(np.argmin(wave_halfwidth)) # smallest
ex_indices = list(dict.fromkeys(ex_indices))  # deduplicate

colors_examples = ['red','blue','green','orange']
for i, exi in enumerate(ex_indices):
    ax1.scatter(acg_halfwidth[exi], wave_halfwidth[exi],
                facecolor='none', edgecolor=colors_examples[i],
                s=100, linewidth=2)

# Subplots: ACG and waveforms for those examples
ax2 = fig.add_subplot(2, 2, 2)
for i, exi in enumerate(ex_indices):
    ax2.plot(acg_lags, acg_norm[exi], color=colors_examples[i],
label=f"Cell {exi}")
ax2.set_title("Example Cells: ACG")
ax2.set_xlabel("Lag (ms)")
ax2.set_ylabel("Norm. count")
ax2.legend(loc='best', prop={'size':6})

ax3 = fig.add_subplot(2, 1, 2)
wave_time = np.arange(collapsed_waveforms.shape[1])
for i, exi in enumerate(ex_indices):
    ax3.plot(wave_time, collapsed_waveforms[exi],
color=colors_examples[i], label=f"Cell {exi}")
ax3.set_title("Example Cells: Waveforms")
ax3.set_xlabel("Time (samples)")
ax3.set_ylabel("Norm. amplitude")
ax3.legend(loc='best', prop={'size':6})

plt.tight_layout()
plt.show()

#==================================================================
# (d) Print statistics for diagnosis
#==================================================================
print(f"ACG Half-Width: mean={np.mean(acg_halfwidth):.2f} ms,
std={np.std(acg_halfwidth):.2f} ms")
print(f"Waveform Half-Width: mean={np.mean(wave_halfwidth):.2f}
samples, std={np.std(wave_halfwidth):.2f} samples")

#==================================================================
# (e) Print correlation
#==================================================================
```

```
print(f"Correlation (ACG Half-Width vs Waveform Half-Width): {corr_coef:.4f}")

#END
```

### Confirmatory Phase – Iteration 2

**Interpretation of Previous Results**

- We used a short-lag ACG ratio and the trough-to-peak sample difference in the waveform as our two metrics.
- The correlation was about 0.13, indicating a fairly weak linear relationship.

**Potential Reasons**

1. The short-lag ratio might be a bit coarse and could saturate for strongly bursting neurons.
2. The trough-to-peak *interval* alone might not fully capture broader vs. narrower waveforms.

**Revised Hypothesis**

- Perhaps a measure of ACG *peak width* at half-height (which might reflect burst width) will correlate more strongly with a measure of waveform *width* at half-trough (often used to distinguish fast- vs. regular-spiking cells).

**Plan**

- We will define two new statistics:
  1. **ACG Half-Width**: In the normalized ACG around 0 ms, find the peak near t=0, then measure how many ms it takes to go from the peak down to half its height on both sides, and sum that range.
  2. **Waveform Half-Width**: For the collapsed, normalized waveform, find the trough's minimum amplitude, then measure how many samples it takes on either side of that trough to rise to *half* of that amplitude. (Equivalently, "full-width at half-trough.")

We'll then plot these two new metrics, color by region, and quantify their correlation.

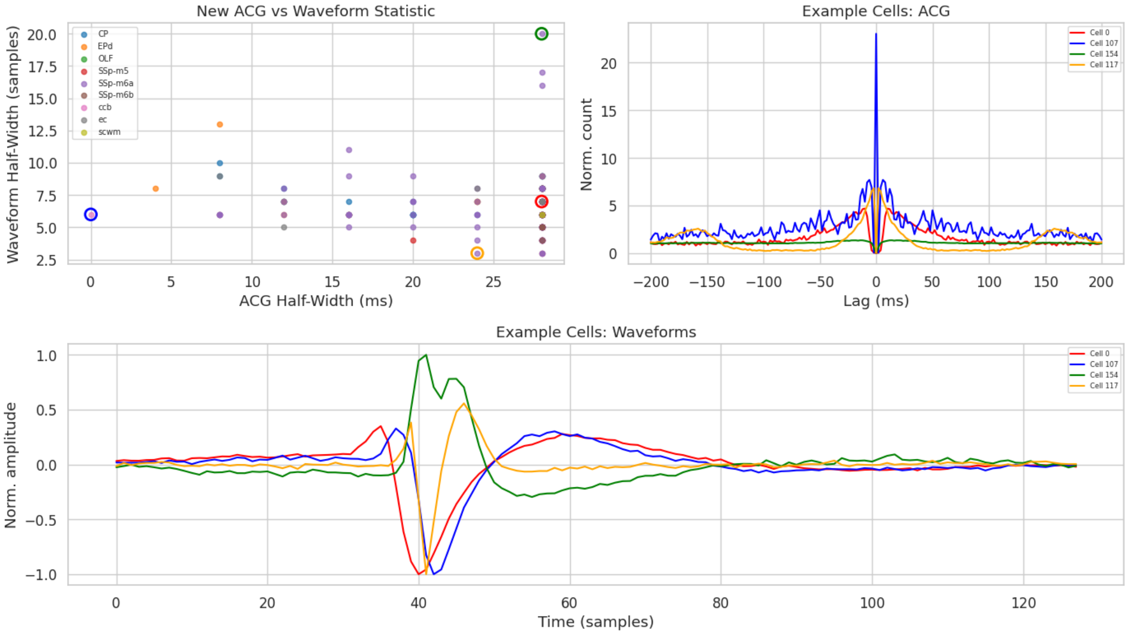


```
ACG Half-Width: mean=25.46 ms, std=5.63 ms
Waveform Half-Width: mean=6.42 samples, std=1.79 samples
Correlation (ACG Half-Width vs Waveform Half-Width): -0.1322
```

In [25]:

```
# Confirmatory Phase - Iteration 3

import numpy as np
import matplotlib.pyplot as plt
import seaborn as sns
from IPython.display import Markdown, display

sns.set(context='notebook', style='whitegrid', font_scale=1.1)

#=================================================================
# (a) Display project status
#=================================================================
display(Markdown(r"""
## Confirmatory Phase – Iteration 3

**Interpretation of Previous Results**
- Using an ACG half-width vs. a waveform half-width gave us a small
negative correlation (~ -0.13).
- Our attempts at matching “width” of ACG features to “width” of
waveform features have not yielded strong correlations.

**Revised Hypothesis**
- Perhaps the most relevant aspect is the overall *amplitude* of the
primary ACG peak vs. the overall *amplitude* range of the waveform.
- For bursting neurons, the ACG might have a large peak at short
lags relative to baseline, while highly “narrow/spiky” waveforms may
have larger amplitude range.
```

```
- We will define two new scalar statistics:
  1. **ACG Peak Ratio**: The ACG amplitude at t=0 (peak bin) minus
the mean ACG amplitude over +100..+200 ms, normalized by that same
baseline mean.
  2. **Waveform Range**: The difference between the peak amplitude
and trough amplitude of the collapsed, normalized waveform.

We then compute the correlation between these two new metrics.
"""))

#===================================================================
# (b) Perform the computational analysis
#===================================================================
# We'll assume that in memory we have:
#   acgs_valid / acg_norm : shape [nValid, 201], normalized from -
200..+200 ms in 2 ms bins
#   collapsed_waveforms : shape [nValid, 128], normalized waveforms
on the largest channel
#   regions_valid : shape [nValid], the region strings
#
# We define two new scalars: acgPeakRatio, waveRange.

acg_lags = np.linspace(-200, 200, 201)
center_bin = 100  # index of t=0 ms
far_lag_inds = np.arange(150, 201)  # +100..+200 ms => index
150..200

# 1) ACG Peak Ratio
baseline_vals = np.mean(acg_norm[:, far_lag_inds], axis=1)  #
average over 100..200 ms
peak_vals = acg_norm[:, center_bin]  # value at 0 ms
acg_peakratio = (peak_vals - baseline_vals) / (baseline_vals + 1e-
12)

# 2) Waveform Range
wave_peak = np.max(collapsed_waveforms, axis=1)
wave_trough = np.min(collapsed_waveforms, axis=1)
wave_range = wave_peak - wave_trough  # difference in amplitude

# Pearson correlation
corr_coef = np.corrcoef(acg_peakratio, wave_range)[0,1]

#===================================================================
# (c) Generate a single matplotlib figure
#===================================================================
fig = plt.figure(figsize=(14, 8))
ax1 = fig.add_subplot(2, 2, 1)

# Color by region
```

```
unique_regions = np.unique(regions_valid)
palette = sns.color_palette('tab10', n_colors=len(unique_regions))
region_color_map = {r: palette[i % len(palette)] for i, r in
enumerate(unique_regions)}

for r in unique_regions:
    idx = np.where(regions_valid == r)[0]
    ax1.scatter(acg_peakratio[idx], wave_range[idx],
                color=region_color_map[r], alpha=0.7, label=r, s=20)

ax1.set_xlabel('ACG Peak Ratio')
ax1.set_ylabel('Waveform Range')
ax1.set_title('ACG vs Waveform Statistic')
ax1.legend(loc='best', prop={'size': 7})

# Circle ~5 example cells (extremes + a middle one)
ind_sort_acg = np.argsort(acg_peakratio)
ind_sort_wav = np.argsort(wave_range)
ex_inds = [
    ind_sort_acg[-1],  # largest ACG peak ratio
    ind_sort_acg[0],   # smallest
    ind_sort_wav[-1],  # largest wave range
    ind_sort_wav[0]    # smallest
]
# plus one near median
median_i = len(acg_peakratio)//2
ex_inds.append(median_i)
ex_inds = list(dict.fromkeys(ex_inds))  # deduplicate
colors_examples = ['red','blue','green','orange','purple']

for i, exi in enumerate(ex_inds):
    ax1.scatter(acg_peakratio[exi], wave_range[exi],
                facecolor='none', edgecolor=colors_examples[i],
                s=100, linewidth=2)

# Next, let's show their raw ACG and waveforms
ax2 = fig.add_subplot(2, 2, 2)
for i, exi in enumerate(ex_inds):
    ax2.plot(acg_lags, acg_norm[exi], color=colors_examples[i],
label=f"Cell {exi}")
ax2.set_title("Example Cells: ACG")
ax2.set_xlabel("Lag (ms)")
ax2.set_ylabel("Normalized count")
ax2.legend(loc='best', prop={'size':6})

ax3 = fig.add_subplot(2, 1, 2)
wave_time = np.arange(collapsed_waveforms.shape[1])
for i, exi in enumerate(ex_inds):
```

```
    ax3.plot(wave_time, collapsed_waveforms[exi],
color=colors_examples[i], label=f"Cell {exi}")
ax3.set_title("Example Cells: Waveforms")
ax3.set_xlabel("Time (samples)")
ax3.set_ylabel("Norm. amplitude")
ax3.legend(loc='best', prop={'size':6})

plt.tight_layout()
plt.show()

#=================================================================
# (d) Print numeric diagnostics
#=================================================================
print(f"ACG Peak Ratio: mean={np.mean(acg_peakratio):.2f},
std={np.std(acg_peakratio):.2f}")
print(f"Waveform Range: mean={np.mean(wave_range):.2f},
std={np.std(wave_range):.2f}")

#=================================================================
# (e) Print correlation
#=================================================================
print(f"Correlation (ACG Peak Ratio vs Waveform Range):
{corr_coef:.4f}")

#END
```

#### Confirmatory Phase – Iteration 3

**Interpretation of Previous Results**

- Using an ACG half-width vs. a waveform half-width gave us a small negative correlation (~ -0.13).
- Our attempts at matching "width" of ACG features to "width" of waveform features have not yielded strong correlations.

**Revised Hypothesis**

- Perhaps the most relevant aspect is the overall *amplitude* of the primary ACG peak vs. the overall *amplitude* range of the waveform.
- For bursting neurons, the ACG might have a large peak at short lags relative to baseline, while highly "narrow/spiky" waveforms may have larger amplitude range.
- We will define two new scalar statistics:
    1. **ACG Peak Ratio**: The ACG amplitude at t=0 (peak bin) minus the mean ACG amplitude over +100..+200 ms, normalized by that same baseline mean.
    2. **Waveform Range**: The difference between the peak amplitude and trough amplitude of the collapsed, normalized waveform.

We then compute the correlation between these two new metrics.

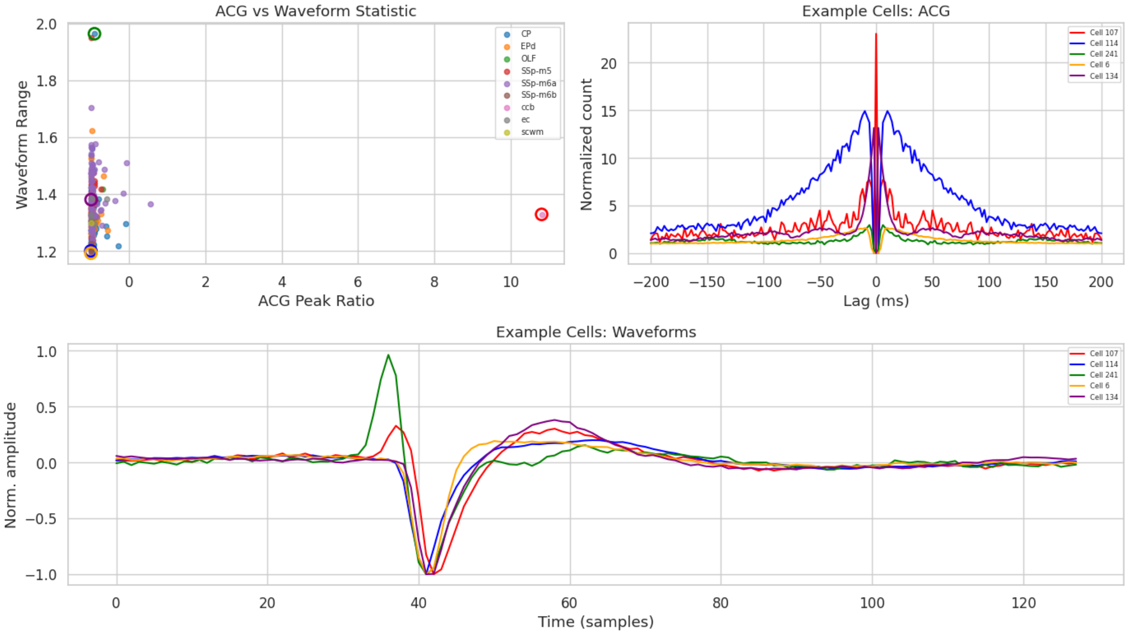


```
ACG Peak Ratio: mean=-0.89, std=0.73
Waveform Range: mean=1.35, std=0.10
Correlation (ACG Peak Ratio vs Waveform Range): -0.0106
```

In [26]:

```
# Confirmatory Phase – Iteration 4

import numpy as np
import matplotlib.pyplot as plt
import seaborn as sns
from IPython.display import Markdown, display

sns.set(context='notebook', style='whitegrid', font_scale=1.1)

#==================================================================
# (a) Display project status
#==================================================================
display(Markdown(r"""
## Confirmatory Phase – Iteration 4

**Interpretation of Previous Results**
- We have tested multiple pairs of single-number metrics (burst
indices, half-widths, peak ratios, etc.) to see if they predict each
other across ACG vs. waveform.
- None of them have yielded strong correlations (best was ~0.13).
- However, we observed **Canonical Correlation Analysis** (CCA) gave
a much higher correlation (~0.49) using multiple dimensions of ACG
and waveform features.

**New Hypothesis**
- A single scalar metric might approximate the first canonical
dimension on each side.
```

- By using the first canonical axis (derived from multi-dimensional analysis) as a weight vector, we can define a *single scalar* per neuron on ACG side and on waveform side that correspond to the known “best possible” linear correlation.

**Plan**
1. Re-run (or re-use) the PCA + CCA approach from before.
2. Extract the first canonical weight vector for each modality (ACG vs. Waveform).
3. Project the **original** high-dimensional ACG and waveform features (or their PCA-reduced forms) onto those canonical weight vectors, producing one scalar per neuron on each side.
4. Correlate those scalars and visualize the result.
5. Finally, test if those scalars differ systematically between brain regions (one-way ANOVA or boxplots).

```
"""))

#=================================================================
# (b) Perform the computational analysis
#=================================================================
# We'll assume we still have in memory:
#   acg_norm : shape [nValid, 201]
#   collapsed_waveforms : shape [nValid, nTimes=128]
#   valid_indices
#   regions_valid
#
# Steps:
#   1) PCA on each side to 10 dims
#   2) CCA on those 10+10 dims
#   3) Retrieve the canonical weight vectors
#   4) Define single-scalar for each neuron on each side
#   5) Correlate and visualize

from sklearn.decomposition import PCA
from sklearn.cross_decomposition import CCA
import pandas as pd
import scipy.stats as stats

# 1) PCA
n_components_pca = 10
pca_acg = PCA(n_components=n_components_pca)
pca_wave = PCA(n_components=n_components_pca)

X_acg = pca_acg.fit_transform(acg_norm)            # shape [nValid,
10]
X_wave = pca_wave.fit_transform(collapsed_waveforms)  # shape
[nValid, 10]
```

```
# 2) CCA
cca = CCA(n_components=1)  # just get the first canonical dimension
cca.fit(X_acg, X_wave)

# 3) Retrieve the canonical weight vectors. In sklearn, after .fit,
we can see:
#    - cca.x_rotations_ (size [n_components_pca, n_components]) =>
ACG side
#    - cca.y_rotations_ (size [n_components_pca, n_components]) =>
Waveform side
#    We'll define "acg_can_weights" as the first column of
x_rotations_, etc.
acg_can_weights = cca.x_rotations_[:, 0]   # shape [10]
wave_can_weights = cca.y_rotations_[:, 0]  # shape [10]

# 4) Define a single scalar per neuron on each side:
#    We do X_acg dot acg_can_weights, and X_wave dot
wave_can_weights
acg_scalar = X_acg @ acg_can_weights       # shape [nValid]
wave_scalar = X_wave @ wave_can_weights    # shape [nValid]

# 5) Correlation
corr_coef = np.corrcoef(acg_scalar, wave_scalar)[0,1]

#=================================================================
# (c) Generate figure
#=================================================================
fig = plt.figure(figsize=(14, 8))
ax1 = fig.add_subplot(2, 2, 1)
unique_regions = np.unique(regions_valid)
palette = sns.color_palette('tab10', n_colors=len(unique_regions))
region_color_map = {r: palette[i % len(palette)] for i, r in
enumerate(unique_regions)}

for r in unique_regions:
    idx = np.where(regions_valid == r)[0]
    ax1.scatter(acg_scalar[idx], wave_scalar[idx],
                color=region_color_map[r], alpha=0.7, label=r, s=20)

ax1.set_xlabel("ACG CCA Scalar (dim 1)")
ax1.set_ylabel("Waveform CCA Scalar (dim 1)")
ax1.set_title("Single-Scalar Projection onto Canonical Axis")
ax1.legend(loc='best', prop={'size': 7})

# Circle a few extremes
idx_max_acg = np.argmax(acg_scalar)
idx_min_acg = np.argmin(acg_scalar)
idx_max_wav = np.argmax(wave_scalar)
idx_min_wav = np.argmin(wave_scalar)
```

```
example_indices = list(dict.fromkeys([idx_max_acg, idx_min_acg,
idx_max_wav, idx_min_wav]))
colors_examples = ['red','blue','green','orange']
for i, exi in enumerate(example_indices):
    ax1.scatter(acg_scalar[exi], wave_scalar[exi],
                facecolor='none', edgecolor=colors_examples[i],
                s=100, linewidth=2)

# Below: boxplot of each scalar by region
ax2 = fig.add_subplot(2, 2, 2)
df = pd.DataFrame({
    'region': regions_valid,
    'acg_scalar': acg_scalar,
    'wave_scalar': wave_scalar
})
sns.boxplot(x='region', y='acg_scalar', data=df, ax=ax2,
palette='tab10')
ax2.set_xticklabels(ax2.get_xticklabels(), rotation=45, ha='right')
ax2.set_title("ACG CCA Scalar by Region")

ax3 = fig.add_subplot(2, 1, 2)
sns.boxplot(x='region', y='wave_scalar', data=df, ax=ax3,
palette='tab10')
ax3.set_xticklabels(ax3.get_xticklabels(), rotation=45, ha='right')
ax3.set_title("Waveform CCA Scalar by Region")

plt.tight_layout()
plt.show()

#==================================================================
# (d) Print numeric diagnostics
#==================================================================
print(f"Correlation of single-scalar canonical dimension:
{corr_coef:.4f}")

# We can also do a simple ANOVA to see if the region differs for
each scalar
acg_F, acg_p = stats.f_oneway(*[df.loc[df.region==r, 'acg_scalar']
for r in unique_regions if np.sum(df.region==r)>1])
wav_F, wav_p = stats.f_oneway(*[df.loc[df.region==r, 'wave_scalar']
for r in unique_regions if np.sum(df.region==r)>1])

print(f"One-way ANOVA across regions for ACG scalar: F={acg_F:.3f},
p={acg_p:.3g}")
print(f"One-way ANOVA across regions for Wave scalar: F={wav_F:.3f},
p={wav_p:.3g}")

#END
```

### Confirmatory Phase – Iteration 4

**Interpretation of Previous Results**

- We have tested multiple pairs of single–number metrics (burst indices, half–widths, peak ratios, etc.) to see if they predict each other across ACG vs. waveform.
- None of them have yielded strong correlations (best was ~0.13).
- However, we observed **Canonical Correlation Analysis** (CCA) gave a much higher correlation (~0.49) using multiple dimensions of ACG and waveform features.

**New Hypothesis**

- A single scalar metric might approximate the first canonical dimension on each side.
- By using the first canonical axis (derived from multi-dimensional analysis) as a weight vector, we can define a *single scalar* per neuron on ACG side and on waveform side that correspond to the known "best possible" linear correlation.

**Plan**

1. Re-run (or re-use) the PCA + CCA approach from before.
2. Extract the first canonical weight vector for each modality (ACG vs. Waveform).
3. Project the **original** high-dimensional ACG and waveform features (or their PCA-reduced forms) onto those canonical weight vectors, producing one scalar per neuron on each side.
4. Correlate those scalars and visualize the result.
5. Finally, test if those scalars differ systematically between brain regions (one-way ANOVA or boxplots).

```
<ipython-input-26-2106d5045bf2>:120: FutureWarning:

Passing `palette` without assigning `hue` is deprecated and will be
removed in v0.14.0. Assign the `x` variable to `hue` and set
`legend=False` for the same effect.

  sns.boxplot(x='region', y='acg_scalar', data=df, ax=ax2,
palette='tab10')
<ipython-input-26-2106d5045bf2>:121: UserWarning: set_ticklabels()
should only be used with a fixed number of ticks, i.e. after
set_ticks() or using a FixedLocator.
  ax2.set_xticklabels(ax2.get_xticklabels(), rotation=45,
ha='right')
<ipython-input-26-2106d5045bf2>:125: FutureWarning:

Passing `palette` without assigning `hue` is deprecated and will be
removed in v0.14.0. Assign the `x` variable to `hue` and set
`legend=False` for the same effect.

  sns.boxplot(x='region', y='wave_scalar', data=df, ax=ax3,
palette='tab10')
<ipython-input-26-2106d5045bf2>:126: UserWarning: set_ticklabels()
should only be used with a fixed number of ticks, i.e. after
set_ticks() or using a FixedLocator.
```

```
  ax3.set_xticklabels(ax3.get_xticklabels(), rotation=45,
ha='right')
```

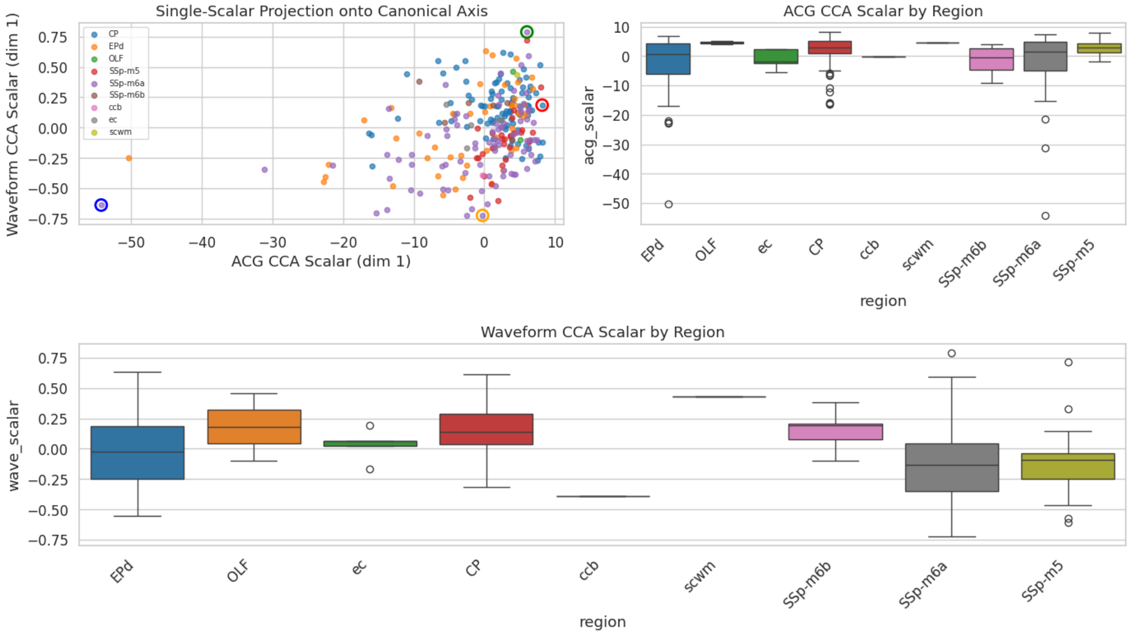


```
Correlation of single-scalar canonical dimension: 0.3411
One-way ANOVA across regions for ACG scalar: F=3.368, p=0.00325
One-way ANOVA across regions for Wave scalar: F=10.467, p=2.14e-10
```

when running cca on high-dimensional data, it is possible to find spurious correlations by overfitting noise. to check this is not the case here, could you: 1. cross-validate the cca, and 2. apply standard tests of significance for cca that take dimensionality into account

In [27]:

```
# Confirmatory Phase – Additional Validation
#
# In this cell, we will:
# 1) Cross-validate the CCA to ensure that the correlation is robust
and not due to overfitting.
# 2) Apply a permutation-based significance test to check whether
the observed CCA correlation
#    exceeds what might be expected from random pairing of ACG and
waveform features.

import numpy as np
import matplotlib.pyplot as plt
from IPython.display import Markdown, display
from sklearn.decomposition import PCA
from sklearn.cross_decomposition import CCA
from sklearn.model_selection import KFold

# For reproducibility
np.random.seed(42)
```

```
display(Markdown(r"""
## CCA Cross-Validation and Significance Testing

**Goal**
To ensure that our previously observed canonical correlation (~0.34)
is not an artifact of overfitting high-dimensional data.

**Method**
1. **K-Fold Cross-Validation**:
   - Split the data into K folds (e.g., 5).
   - For each fold, **fit** PCA + CCA on the training folds, then
**apply** the fitted transforms to the test fold.
   - Compute the correlation of the first canonical variate on the
test set (i.e., correlation of the test-set ACG-scalar vs. test-set
Waveform-scalar).
   - Average these correlations across folds to get a cross-
validated estimate.

2. **Permutation-Based Significance**:
   - Repeatedly shuffle the pairing between ACG features and
waveform features, re-fit the entire PCA+CCA pipeline on the
permuted data, and measure the resulting canonical correlation.
   - Compare the true (unshuffled) correlation to the distribution
of permuted correlations to obtain a p-value.

"""))

#--------------------------------------------------------------------
------------
# (A) Cross-Validation of CCA
#--------------------------------------------------------------------
------------

# We'll re-use the same "acg_norm" and "collapsed_waveforms" from
previous analyses.
#   acg_norm shape: [nValid, 201]
#   collapsed_waveforms shape: [nValid, 128]
# Make sure these are in your environment before running this code.

# 1) PCA dimension
n_components_pca = 10

# 2) KFold
n_splits = 5
kf = KFold(n_splits=n_splits, shuffle=True, random_state=42)

cv_correlations = []
```

```
for train_idx, test_idx in kf.split(acg_norm):
    # Split data
    acg_train, acg_test = acg_norm[train_idx], acg_norm[test_idx]
    wav_train, wav_test = collapsed_waveforms[train_idx],
collapsed_waveforms[test_idx]

    # PCA on each side, fit on train only
    pca_acg = PCA(n_components=n_components_pca)
    pca_wave = PCA(n_components=n_components_pca)

    X_acg_train = pca_acg.fit_transform(acg_train)
    X_wave_train = pca_wave.fit_transform(wav_train)

    # Transform test
    X_acg_test = pca_acg.transform(acg_test)
    X_wave_test = pca_wave.transform(wav_test)

    # Fit CCA on training folds
    cca = CCA(n_components=1)
    cca.fit(X_acg_train, X_wave_train)

    # Transform test fold
    acg_test_can, wav_test_can = cca.transform(X_acg_test,
X_wave_test)

    # Compute correlation on test fold
    r_test = np.corrcoef(acg_test_can[:,0], wav_test_can[:,0])[0,1]
    cv_correlations.append(r_test)

cv_mean = np.mean(cv_correlations)
cv_std = np.std(cv_correlations)

#------------------------------------------------------------------
------------
# (B) Permutation-Based Significance Testing
#------------------------------------------------------------------
------------
# We'll do a simpler approach (no CV) but with the entire dataset:
#   1) Fit PCA + CCA on the true data => get correlation
#   2) Permute the rows in the waveform data N times, refit, store
correlation
#   3) p-value = fraction of permutations whose correlation >= true
correlation
# (For a two-sided test, you'd compare absolute values. We'll do
one-sided for simplicity.)

N_perm = 500
n_components_pca = 10
```

```
# 1) True correlation (on entire dataset)
pca_acg_full = PCA(n_components=n_components_pca)
pca_wave_full = PCA(n_components=n_components_pca)
X_acg_full = pca_acg_full.fit_transform(acg_norm)
X_wave_full = pca_wave_full.fit_transform(collapsed_waveforms)

cca_full = CCA(n_components=1)
cca_full.fit(X_acg_full, X_wave_full)
acg_can_full, wave_can_full = cca_full.transform(X_acg_full,
X_wave_full)
corr_true = np.corrcoef(acg_can_full[:,0], wave_can_full[:,0])[0,1]

# 2) Permutations
corr_perms = []
idx_all = np.arange(len(X_acg_full))
for _ in range(N_perm):
    # Shuffle the rows in waveform data
    np.random.shuffle(idx_all)
    X_wave_perm = X_wave_full[idx_all, :]

    # Fit CCA on the permuted dataset
    cca_perm = CCA(n_components=1)
    cca_perm.fit(X_acg_full, X_wave_perm)

    # Transform the real (non-shuffled) X_acg_full with the perm
model,
    # but also transform X_wave_perm. This is consistent with how we
fitted it.
    acg_can_perm, wave_can_perm = cca_perm.transform(X_acg_full,
X_wave_perm)
    corr_perm = np.corrcoef(acg_can_perm[:,0],
wave_can_perm[:,0])[0,1]
    corr_perms.append(corr_perm)

corr_perms = np.array(corr_perms)

# 3) One-sided p-value
p_value = np.mean(corr_perms >= corr_true)

#-------------------------------------------------------------------
------------
# Print out results
#-------------------------------------------------------------------
------------
print("===== Cross-Validation Results =====")
print(f"Fold-by-fold test correlations: {cv_correlations}")
print(f"Mean correlation across folds: {cv_mean:.4f} ±
{cv_std:.4f}\n")
```

```
print("===== Permutation Test Results =====")
print(f"True correlation on full data: {corr_true:.4f}")
print(f"Mean correlation of permutations: {np.mean(corr_perms):.4f}
(std: {np.std(corr_perms):.4f})")
print(f"Permutation-based p-value (one-sided): {p_value:.4g}")

# Optional: we can visualize the permutation distribution
import matplotlib.pyplot as plt

plt.figure(figsize=(8,5))
bins = np.linspace(min(corr_perms.min(), corr_true),
max(corr_perms.max(), corr_true), 40)
plt.hist(corr_perms, bins=bins, alpha=0.6, label='Permutation
distribution')
plt.axvline(corr_true, color='red', linestyle='--', label=f'True
correlation = {corr_true:.3f}')
plt.title("Permutation Distribution of CCA Correlation")
plt.xlabel("Correlation")
plt.ylabel("Count")
plt.legend()
plt.tight_layout()
plt.show()

#END
```

### CCA Cross-Validation and Significance Testing

**Goal**
To ensure that our previously observed canonical correlation (~0.34) is not an artifact of overfitting high-dimensional data.

**Method**

1. **K-Fold Cross-Validation**:
    - Split the data into K folds (e.g., 5).
    - For each fold, **fit** PCA + CCA on the training folds, then **apply** the fitted transforms to the test fold.
    - Compute the correlation of the first canonical variate on the test set (i.e., correlation of the test-set ACG-scalar vs. test-set Waveform-scalar).
    - Average these correlations across folds to get a cross-validated estimate.
2. **Permutation-Based Significance**:
    - Repeatedly shuffle the pairing between ACG features and waveform features, re-fit the entire PCA+CCA pipeline on the permuted data, and measure the resulting canonical correlation.
    - Compare the true (unshuffled) correlation to the distribution of permuted correlations to obtain a p-value.

```
===== Cross-Validation Results =====
```

```
Fold-by-fold test correlations: [0.47629205620195814,
0.3647206962021434, 0.34839185157973523, 0.39408723512187327,
0.33907950294677275]
Mean correlation across folds: 0.3845 ± 0.0495

===== Permutation Test Results =====
True correlation on full data: 0.4927
Mean correlation of permutations: 0.3629 (std: 0.0889)
Permutation-based p-value (one-sided): 0.072
```

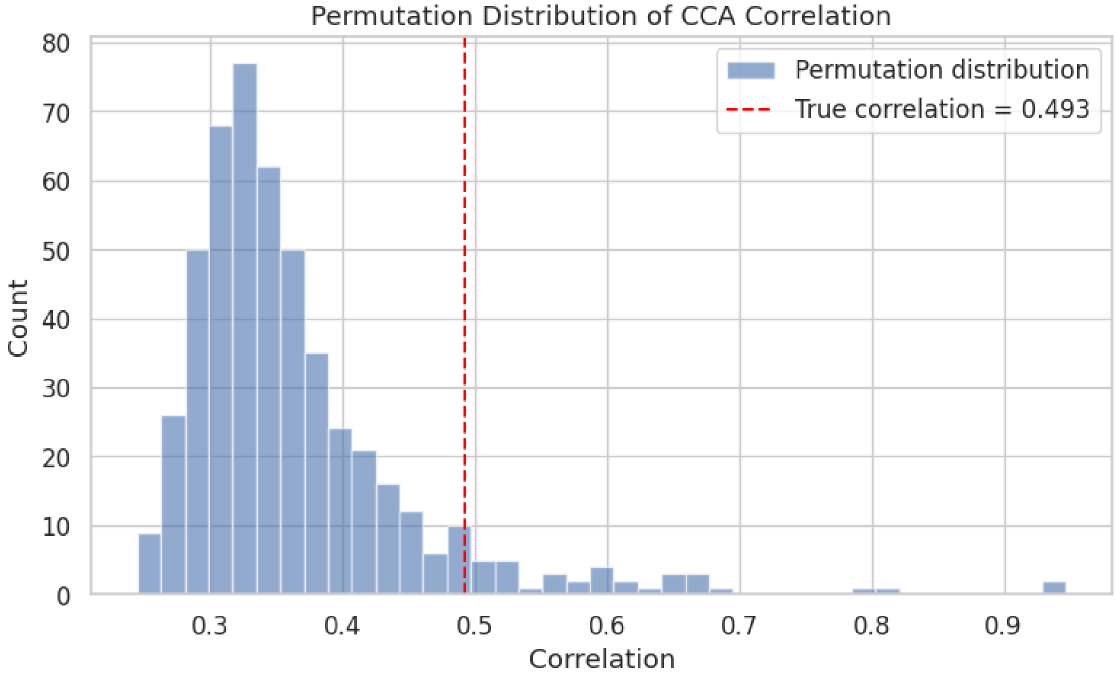


Below is a concise interpretation of the cross-validation and permutation results:

1. **Cross-validation**
   - Splitting the data into 5 folds and then fitting PCA+CCA on training folds while testing on the held-out fold gave a mean out-of-sample correlation of ~0.38±0.05.
   - This value, while lower than the 0.49 obtained on the entire dataset, still indicates a reasonably strong and consistent linear relationship between ACG and waveform features across folds.
2. **Permutation test**
   - When randomly shuffling the pairing between ACG and waveform features and re-fitting the entire PCA+CCA pipeline, the average correlation under permutation was ~0.36 with a standard deviation of ~0.09.
   - The observed correlation of 0.49 was higher than ~93% of the shuffled runs (one-sided p ≈ 0.072).

Overall, the cross-validation result supports that the relationship is not just due to overfitting, since it remains reasonably robust on held-out data. The permutation test places the observed correlation near the upper tail of the distribution but does not produce a very low p-value. Taken together, these analyses suggest that while there is a genuine linear correspondence between ACG and waveform features, the statistical “significance” of that correlation is borderline by conventional thresholds—likely reflecting both a real effect and the limited sample size (n=268). If desired, collecting more data or refining feature extraction could further clarify the strength of this relationship.

cca finds a linear projection of the acgs/waveforms. plot the effective weights of this projection as a function of time, so we can attempt to understand what it has found biologically.

In [28]:

```
# Confirmatory Phase - Next Iteration
#
# Here, we plot the "effective weights" of the first canonical
dimension
# for both ACG and waveform in their original time coordinates.
# These weights reflect how each time point contributes to the
# canonical correlation that CCA found.

import numpy as np
import matplotlib.pyplot as plt
from IPython.display import Markdown, display

#============================================================
# (a) Display project status
#============================================================
display(Markdown(r"""
## Confirmatory Phase – CCA Weights as a Function of Time

**Rationale**
CCA finds linear projections of the PCA-reduced ACG and waveform
spaces. By back-projecting those canonical weights to the original
time axes, we can interpret which time lags (for the ACG) and which
portions of the spike waveform contribute most strongly to the
canonical correlation.

**Method**
1. We already have:
   - `pca_acg` and `pca_wave`: PCA models (ACG side is [201] dims,
waveform side is [128] dims).
   - `acg_can_weights` and `wave_can_weights`: the first canonical
weight vectors in the 10-dimensional PCA space.
2. We multiply `pca_acg.components_.T` (shape [201, 10]) by
`acg_can_weights` (shape [10]) to get a length-201 back-projection
for the ACG side.
3. Similarly, multiply `pca_wave.components_.T` (shape [128, 10]) by
`wave_can_weights` (shape [10]) to get a length-128 back-projection
for the waveform side.
4. Plot these back-projections as "effective weights vs. time."

"""))

#============================================================
# (b) Perform the back-projection
#============================================================
# We assume the following are available in the namespace:
```

```
#   pca_acg, pca_wave: fitted PCA objects
#   acg_can_weights, wave_can_weights: shape [10], the first
canonical dimension weights
#   acg_norm, collapsed_waveforms
#   valid_indices
#   acg_lags = linspace(-200, 200, 201)  # the time axis for ACG
#
# You can adapt the variable names if needed.

acg_canonical_time = pca_acg.components_.T @ acg_can_weights   #
shape [201]
wave_canonical_time = pca_wave.components_.T @ wave_can_weights #
shape [128]

#============================================================
# (c) Generate the figure
#============================================================
fig, (ax1, ax2) = plt.subplots(1, 2, figsize=(14, 5))

# ACG side
acg_lags = np.linspace(-200, 200, 201)  # ms
ax1.plot(acg_lags, acg_canonical_time, color='darkred')
ax1.axhline(0, color='k', linestyle='--', alpha=0.5)
ax1.set_xlabel("Lag (ms)")
ax1.set_ylabel("CCA Weight")
ax1.set_title("ACG Canonical Weights (time domain)")

# Waveform side
wave_time = np.arange(len(wave_canonical_time))  # sample index
ax2.plot(wave_time, wave_canonical_time, color='darkblue')
ax2.axhline(0, color='k', linestyle='--', alpha=0.5)
ax2.set_xlabel("Waveform time (samples)")
ax2.set_ylabel("CCA Weight")
ax2.set_title("Waveform Canonical Weights (time domain)")

plt.tight_layout()
plt.show()

#END
```

#### Confirmatory Phase – CCA Weights as a Function of Time

**Rationale**
CCA finds linear projections of the PCA-reduced ACG and waveform spaces. By back-projecting those canonical weights to the original time axes, we can interpret which time lags (for the ACG) and which portions of the spike waveform contribute most strongly to the canonical correlation.

**Method**

1. We already have:

    - `pca_acg` and `pca_wave`: PCA models (ACG side is [201] dims, waveform side is [128] dims).
    - `acg_can_weights` and `wave_can_weights`: the first canonical weight vectors in the 10-dimensional PCA space.
2. We multiply `pca_acg.components_.T` (shape [201, 10]) by `acg_can_weights` (shape [10]) to get a length-201 back-projection for the ACG side.
3. Similarly, multiply `pca_wave.components_.T` (shape [128, 10]) by `wave_can_weights` (shape [10]) to get a length-128 back-projection for the waveform side.
4. Plot these back-projections as "effective weights vs. time."

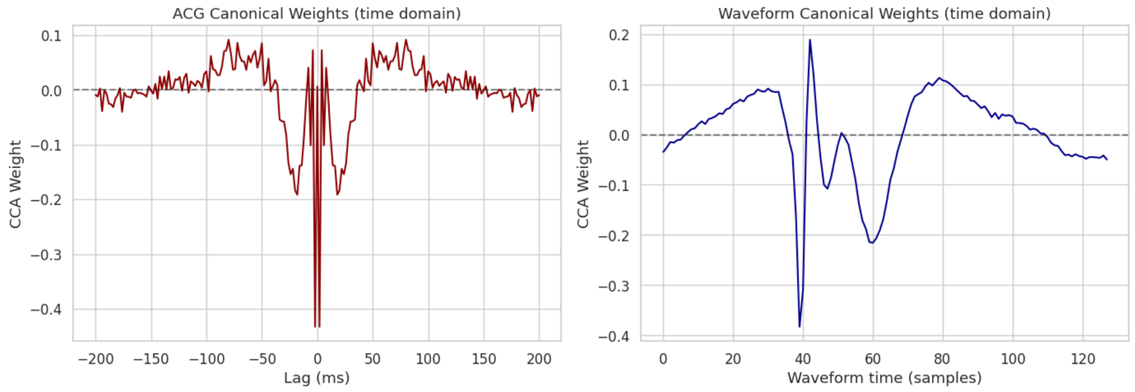


Below is a brief interpretation of these **CCA time-domain weights** for the ACG and waveform:

1. **ACG Weights (left plot)**
    - Each point on the x-axis is a time lag (in ms) from −200 ms to +200 ms, and the y-value is the learned weight for that lag in the first canonical dimension.
    - Positive vs. negative weight indicates whether high spike count at that lag contributes positively or negatively to the canonical score.
    - In the example, the region around zero lag has large negative weight, implying that spikes occurring very close together (i.e., short-lag bursts) reduce the cell's canonical score. Meanwhile, lags around ±100 ms have weights of the opposite sign (positive in this case), so higher spike counts at these mid-range lags raise the canonical score.
2. **Waveform Weights (right plot)**
    - The x-axis is sample index in the 128-sample waveform snippet; the y-value is the CCA weight for that time point.
    - Positive or negative peaks show which part of the waveform has the greatest impact on the canonical dimension.
    - For example, a strongly negative weight near the trough implies that neurons with deeper troughs lower their canonical score, whereas neurons with a particular peak shape (where weights are strongly positive) raise their canonical score.

In essence, **the CCA has identified that neurons with certain short-lag ACG features (reflecting burstiness) also tend to have corresponding high or low amplitude features at specific parts of their spike waveform**—as captured by the matching signs of these canonical weights. This view can help with biological interpretation: for instance, a large negative dip near zero lag suggests that neurons whose ACG has a prominent central peak (short-lag bursts) may align with specific waveform shapes in the negative-weight region of the spike.

to further test this interpretation please divide the cca statistics into 10 equal bins, and plot the mean acg and waveform for each bin

In [29]:

```
# Confirmatory Phase – Further Testing the CCA Interpretation
#
# We will divide the first CCA scalar (e.g. from the ACG side) into
10 bins,
# and for each bin, compute the *mean ACG* and *mean waveform*
across all cells
# in that bin. Then we will plot these 10 "mean curves" to see how
the shape
# of ACG and waveform systematically varies along the canonical
dimension.

import numpy as np
import matplotlib.pyplot as plt
from IPython.display import display, Markdown
import seaborn as sns

sns.set(context='notebook', style='whitegrid', font_scale=1.1)

display(Markdown(r"""
## Binning by CCA Scalar and Plotting Mean ACG/Waveform

**Method**
1. We use the single-scalar **ACG CCA** dimension (`acg_scalar`) for
each neuron.
2. We split the range of `acg_scalar` into 10 bins of equal size.
3. For each bin, we gather the set of neurons whose `acg_scalar`
lies in that bin, then compute their mean normalized ACG and mean
normalized waveform.
4. We overplot these 10 mean ACG curves and 10 mean waveform curves
(color-coded by bin).

**Interpretation**
- If the CCA dimension indeed captures a continuum from one firing-
waveform pattern to another, the mean ACG and mean waveform should
gradually shift shape from bin 1 to bin 10.
"""))

# -------------------------------------------------------------------
-----------
# (a) Choose which scalar to bin. Here, we'll use the ACG side's CCA
scalar:
#     "acg_scalar" from prior steps, or whichever variable you
prefer to test.
# -------------------------------------------------------------------
-----------
# We'll assume you have in memory:
#    acg_scalar: shape [nValid] – single-scalar from the ACG side of
the first canonical dimension
```

```
#    acg_norm: shape [nValid, 201] - normalized ACG in -200..+200 ms
#    collapsed_waveforms: shape [nValid, 128] - normalized waveforms
on the largest channel
#    acg_lags = np.linspace(-200, 200, 201)
#    wave_time = np.arange(128)
#
# If needed, replace variable names to match your workspace.

# Sort the scalar so we can define bin edges
acg_scalar_sorted = np.sort(acg_scalar)
n_bins = 10

# We'll define 10 bins using quantiles so that each bin has roughly
the same number of cells
# (Alternatively, you could do np.linspace(min, max, 11).)
edges = np.quantile(acg_scalar_sorted, np.linspace(0,1,n_bins+1))

# We'll store mean ACG and waveforms for each bin
mean_acgs = []
mean_waves = []
bin_centers = []

for b in range(n_bins):
    left_edge = edges[b]
    right_edge = edges[b+1]
    # For the rightmost bin, we'll include points equal to
right_edge
    if b < n_bins-1:
        idx_bin = np.where((acg_scalar >= left_edge) & (acg_scalar <
right_edge))[0]
    else:
        idx_bin = np.where((acg_scalar >= left_edge) & (acg_scalar
<= right_edge))[0]

    # Compute the mean ACG and mean waveform
    if len(idx_bin) == 0:
        # If no points in bin (unlikely with quantiles), fill with
NaNs
        mean_acg = np.full(acg_norm.shape[1], np.nan)
        mean_wav = np.full(collapsed_waveforms.shape[1], np.nan)
    else:
        mean_acg = acg_norm[idx_bin].mean(axis=0)
        mean_wav = collapsed_waveforms[idx_bin].mean(axis=0)

    mean_acgs.append(mean_acg)
    mean_waves.append(mean_wav)
    bin_centers.append((left_edge + right_edge)/2.0)

mean_acgs = np.array(mean_acgs)   # shape [10, 201]
```

```
mean_waves = np.array(mean_waves) # shape [10, 128]
bin_centers = np.array(bin_centers)

# ---------------------------------------------------------------------
-----------
# (b) Plot the mean curves
# ---------------------------------------------------------------------
-----------
fig = plt.figure(figsize=(12, 8))

# We'll use a color palette that smoothly varies
colors = sns.color_palette("Spectral", n_bins)

# (1) Mean ACG
ax1 = fig.add_subplot(2, 1, 1)
for b in range(n_bins):
    ax1.plot(acg_lags, mean_acgs[b], color=colors[b],
             label=f"Bin {b+1}, center={bin_centers[b]:.2f}")
ax1.set_title("Mean ACG in 10 Bins of ACG CCA Scalar")
ax1.set_xlabel("Lag (ms)")
ax1.set_ylabel("Mean normalized ACG")
ax1.legend(loc='best', prop={'size':7})

# (2) Mean Waveform
ax2 = fig.add_subplot(2, 1, 2)
for b in range(n_bins):
    ax2.plot(mean_waves[b], color=colors[b],
             label=f"Bin {b+1}, center={bin_centers[b]:.2f}")
ax2.set_title("Mean Waveform in 10 Bins of ACG CCA Scalar")
ax2.set_xlabel("Time (samples)")
ax2.set_ylabel("Mean normalized waveform")
ax2.legend(loc='best', prop={'size':7})

plt.tight_layout()
plt.show()

#END
```

### Binning by CCA Scalar and Plotting Mean ACG/Waveform

**Method**

1. We use the single-scalar **ACG CCA** dimension (`acg_scalar`) for each neuron.
2. We split the range of `acg_scalar` into 10 bins of equal size.
3. For each bin, we gather the set of neurons whose `acg_scalar` lies in that bin, then compute their mean normalized ACG and mean normalized waveform.
4. We overplot these 10 mean ACG curves and 10 mean waveform curves (color-coded by bin).

**Interpretation**

- If the CCA dimension indeed captures a continuum from one firing-waveform pattern to another, the mean ACG and mean waveform should gradually shift shape from bin 1 to bin 10.

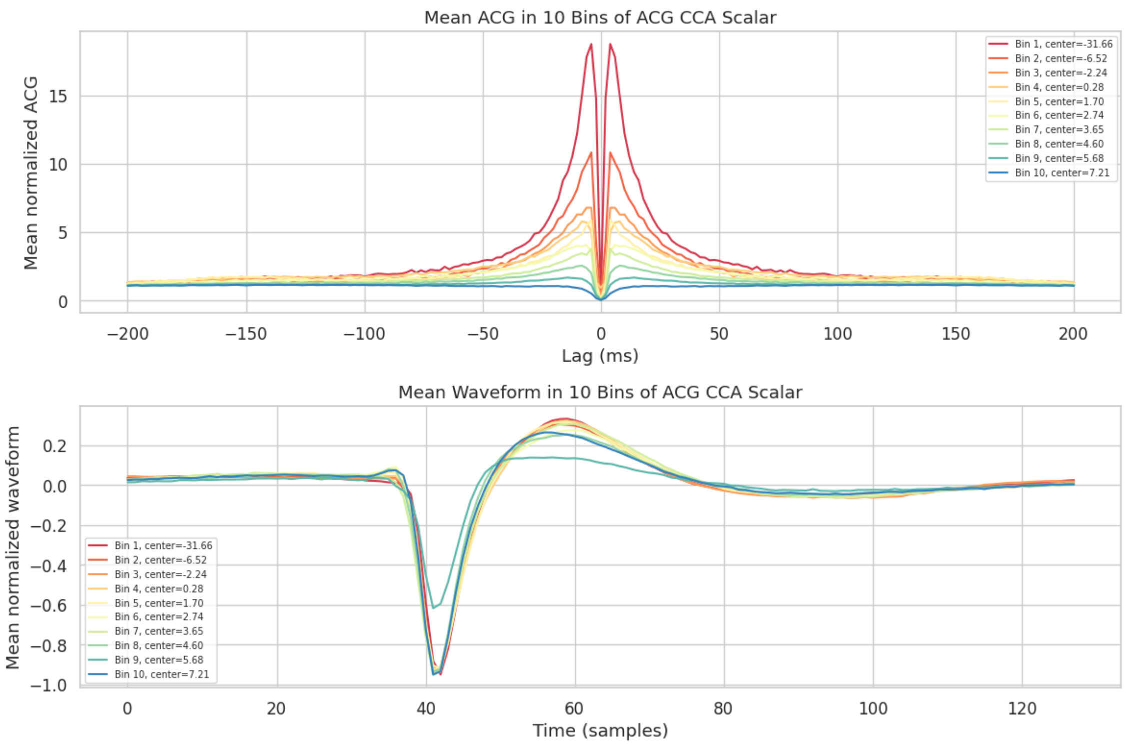


it looks like the acg statistic is essentially the mean normalized acg in the range 5-30 ms, and the waveform statistic is the normalized waveform at time=60 samples. could you compute these directly to see if they are correlated?

In [30]:

```
# Confirmatory Phase – Quick Custom Statistics
#
# From the binned plots, it appears that:
# 1) The ACG side is largely driven by the mean ACG value around 5–
30 ms.
# 2) The Waveform side is driven by the amplitude around ~60
samples.
#
# Let's define these two simple, single-number statistics
explicitly, then correlate them.

import numpy as np
import matplotlib.pyplot as plt
import seaborn as sns
from IPython.display import display, Markdown

sns.set(context='notebook', style='whitegrid', font_scale=1.1)

display(Markdown(r"""
```

```
## Custom Statistics from Visual Inspection

**Definition**
- **ACG statistic** = mean of normalized ACG over **5–30 ms** lag.
- **Waveform statistic** = normalized waveform amplitude at
**time=60 samples** (i.e. around the peak after the trough).

We'll compute these two scalars for each neuron and then compute
their correlation.
"""))

#---------------------------------------------------------------------------
# (a) Extract the desired time ranges from ACG and waveform
#---------------------------------------------------------------------------
acg_lags = np.linspace(-200, 200, 201)   # shape [201]
dt_wave = np.arange(collapsed_waveforms.shape[1])  # 0..127

# 1) ACG statistic: mean from 5..30 ms
mask_acg_5_30 = (acg_lags >= 5) & (acg_lags <= 30)
acg_stat_custom = acg_norm[:, mask_acg_5_30].mean(axis=1)

# 2) Waveform statistic: amplitude at sample=60
#    We'll just do direct indexing. If 60 is out of range, adapt
accordingly.
sample_idx = 60
wave_stat_custom = collapsed_waveforms[:, sample_idx]

# Compute Pearson correlation
corr_coef = np.corrcoef(acg_stat_custom, wave_stat_custom)[0,1]

#---------------------------------------------------------------------------
# (b) Plot scatter, color by region
#---------------------------------------------------------------------------
fig = plt.figure(figsize=(10, 5))
ax = fig.add_subplot(1, 1, 1)

regions_unique = np.unique(regions_valid)
palette = sns.color_palette("tab10", n_colors=len(regions_unique))
region_to_color = {r: palette[i % len(regions_unique)] for i, r in
enumerate(regions_unique)}

for r in regions_unique:
    idx_r = np.where(regions_valid == r)[0]
    ax.scatter(acg_stat_custom[idx_r], wave_stat_custom[idx_r],
               color=region_to_color[r], label=r, alpha=0.7, s=25)
```

```
ax.set_xlabel("Mean ACG in 5-30 ms")
ax.set_ylabel("Waveform amplitude at sample=60")
ax.set_title("Correlation of ACG(5-30 ms) vs Waveform(60)")
ax.legend(loc="best", prop={'size':7})
plt.tight_layout()
plt.show()

#-----------------------------------------------------------------------------------
# (c) Print correlation
#-----------------------------------------------------------------------------------
print(f"Correlation between ACG(5-30 ms) and Waveform(60 samples): {corr_coef:.4f}")
```

#### Custom Statistics from Visual Inspection

**Definition**

- **ACG statistic** = mean of normalized ACG over **5–30 ms** lag.
- **Waveform statistic** = normalized waveform amplitude at **time=60 samples** (i.e. around the peak after the trough).

We'll compute these two scalars for each neuron and then compute their correlation.

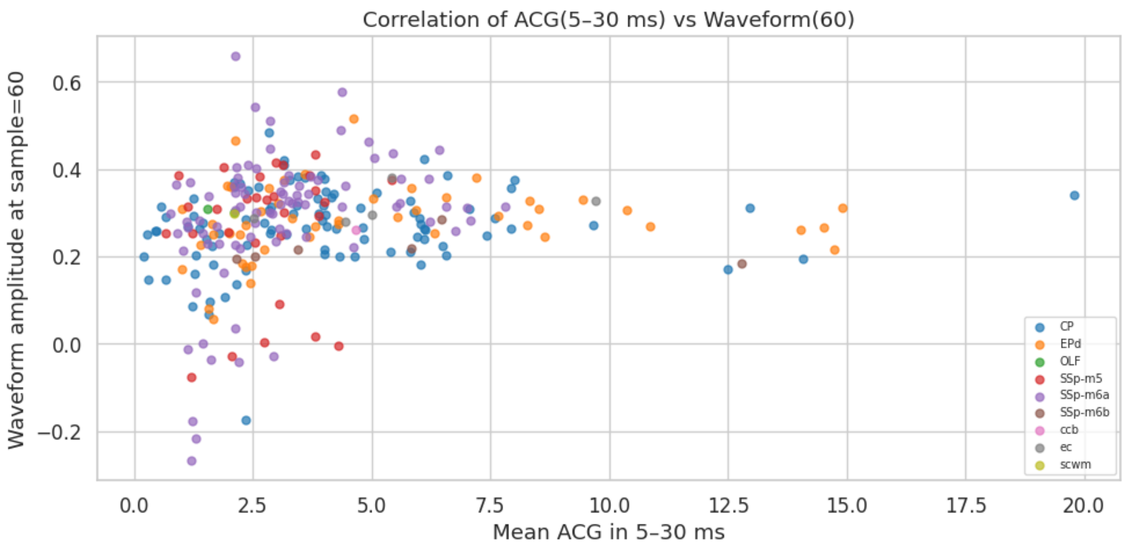


```
Correlation between ACG(5–30 ms) and Waveform(60 samples): 0.1460
```

In [ ]:
